# Generation of a HiBiT-expressing recombinant rat hepacivirus supporting both *in vivo* and *in vitro* infection

**DOI:** 10.1101/2025.11.25.690605

**Authors:** Yasunori Akaike, Tomohisa Tanaka, Hirotake Kasai, Atsuya Yamashita, Yoshiharu Matsuura, Kohji Moriishi

**Affiliations:** Department of Microbiology, Faculty of Medicine, Graduate Faculty of Interdisciplinary Research, University of Yamanashi, Yamanashi 409-3898, Japan; Division of Hepatitis Virology, Institute for Genetic Medicine, Hokkaido University, Hokkaido 060-0815, Japan; Center for Infectious Disease Education and Research, Osaka University, Osaka 565-0871, Japan; Laboratory of Virus Control, Research Institute for Microbial Diseases, Osaka University, Osaka 565-0871, Japan; Center for Life Science Research, University of Yamanashi, Yamanashi 409-3898, Japan

## Abstract

The lack of immuno-competent animal models of hepatitis C virus (HCV) infection has been an obstacle to vaccine development and research on immune responses. *Hepacivirus ratti* (Norway rat hepacivirus-1: NRHV1) is a virus closely related to HCV that specifically infects the liver and induces hepatocellular carcinoma in rats, making it a promising surrogate model for HCV. NRHV1 expressing a reporter gene serves as a powerful tool for analyzing the *in vivo* dynamics and pathogenicity mechanisms of NRHV1. In this study, we developed a reporter NRHV1 capable of infection and replication in both mice and cultured cells and established a platform for generating diverse reporter viruses. A reporter virus containing the HiBiT gene in the coding region of NS5A domain III was constructed using circular polymerase extension reaction (CPER). Infection with this reporter virus led to HiBiT activity in infected cells and the activity was correlated with the amount of intracellular viral RNA. In addition, this reporter virus established persistent infection in NOD–SCID mice and led to the generation of HiBiT activity in the livers of infected mice. Furthermore, reporter virus recovered from infected mice could infect and generate HiBiT activity in cultured cells. These results demonstrate that the reporter virus can infect hepatocytes *in vivo* and *in vitro*. This platform provides a versatile tool for *in vitro* quantitative antiviral screening and *in vivo* pathogenicity studies.

**Author summary:** Hepatitis C virus (HCV) infects millions of people and causes severe liver diseases, yet vaccine development has been hindered by the lack of practical small-animal models with intact immunity. *Hepacivirus ratti* (NRHV1), a close relative of HCV that naturally infects rats, is a promising HCV surrogate, but tools for monitoring NRHV1 infection dynamics have been limited. Here, we developed the first reporter NRHV1 by inserting a small luminescent HiBiT tag into a tolerable region of the NS5A protein. Using a CPER-based reverse genetics system, we generated a recombinant virus that efficiently infected cultured cells and established persistent infection in mice while retaining liver tropism. HiBiT activity closely reflected viral RNA levels, enabling the quantitative monitoring of infection and sensitive evaluation of antiviral drugs and neutralizing antibodies. This reporter NRHV1 system provides a versatile platform for studying hepacivirus pathogenesis and antiviral immunity and will support future efforts toward HCV vaccine development.

## Introduction

Hepatitis C virus (HCV) chronically infects approximately 50.7 million people worldwide, with 1.04 million new cases annually [1]. Chronic infection can cause persistent inflammation, fatty liver, cirrhosis, and hepatocellular carcinoma (HCC). Direct-acting antivirals (DAAs) have greatly improved patient treatment, but no vaccines exist, making patients vulnerable to reinfection [2, 3]. Although animal models are essential for vaccine development, practical immune-competent models are lacking. Chimpanzees and tupaia can be experimentally infected, but they do not fully reproduce the characteristics of human disease, and their research use is limited by ethical and technical considerations [4–6]. No small animal models with intact immune systems can fully recapitulate HCV infection [7, 8]. This limitation hinders research on HCV pathogenesis, immune responses, and vaccine development.

HCV is a member of the *Hepacivirus* genus (*Flaviviridae*), whose relatives infect diverse animals, such as bats, dogs, horses, fish, and rodents, and typically target the liver [9]. HCV itself does not efficiently propagate in nonhuman hosts except for chimpanzees, nor even in mice engineered to express human entry factors. Therefore, related viruses have been explored as alternative models. One example is *Hepacivirus ratti* (Norway rat hepacivirus-1: NRHV1 rn-1 prototype strain), which was isolated from Norway rats [10]. NRHV1 has a 9,656-nt positive-sense RNA genome encoding a polyprotein of 2,958 aa, with organization and processing similar to those of HCV; it shares key features with HCV, including hepatotropism and conserved miR-122 binding sites [11]. NRHV1 causes persistent infection with high-level viremia in rats, immunodeficient mice, and mice whose CD4^+^ cells are transiently depleted before infection, and can induce hepatocellular carcinoma [12]. Thus, NRHV1 provides a valuable model for studying HCV persistence and liver disease.

A recombinant virus encoding a reporter gene is a powerful tool for studying viral dynamics *in vivo*. However, no reporter NRHV1 has yet been described. Here, we identified an appropriate insertion site for introducing the HiBiT gene into the NRHV1 genome and subsequently assessed the infectivity of the reporter virus both *in vitro* and *in vivo*. Our developed reporter virus could infect cultured cells and immune-deficient mice. Furthermore, viral preparations originating from these infected cultured cells or murine sera were infectious *in vitro* and *in vivo*.

## Results

### Preparation of recombinant NRHV1 by CPER

Reverse genetics systems for orthoflaviviruses, SARS-CoV-2, and other viruses have been successfully established using circular polymerase extension reaction (CPER) [13–16]. Therefore, we attempted to develop a method to generate recombinant NRHV1 by CPER for the introduction of an exogenous reporter gene. The McA-RH7777-derived cell line McA1.8 is highly susceptible to a cell-adapted strain of NRHV1 (strain rn-1cp7), which was obtained after seven serial passages in McA1.8 cells, as previously reported [17]. Recombinant NRHV1 was synthesized from the rn-1cp7 strain by CPER as described in the Materials and Methods section (Figure 1A). The circular CPER product was transfected into McA1.8 cells. Intracellular and extracellular levels of NRHV1 RNA were higher in cells transfected with CPER products containing DNA polymerase than in those transfected with control preparations lacking DNA polymerase (Figure 1B). The McA1.8/mCnls-MAVS cell line, which has been reported as an indicator cell line for NRHV1 infection [17], was used to assess infectivity. This cell line expresses a recombinant mCherry protein fused with a nuclear localization signal and an NRHV1-dependent MAVS cleavage site, resulting in the nuclear translocation of mCherry upon NRHV1 infection [17]. Cells transfected with CPER preparations containing or lacking DNA polymerase were harvested 15 days post-transfection (dpt), and their supernatants were inoculated into McA1.8/mCnls-MAVS cells. In cells inoculated with preparations containing DNA polymerase, mCherry fluorescence translocated to the nucleus, whereas it remained cytoplasmic in cells inoculated with preparations lacking DNA polymerase (Figure 1C, left panel). Similarly, intracellular viral RNA levels were higher in cells treated with preparations containing DNA polymerase than in those treated with preparations lacking DNA polymerase (Figure 1C, right panel).

**Figure 1.**
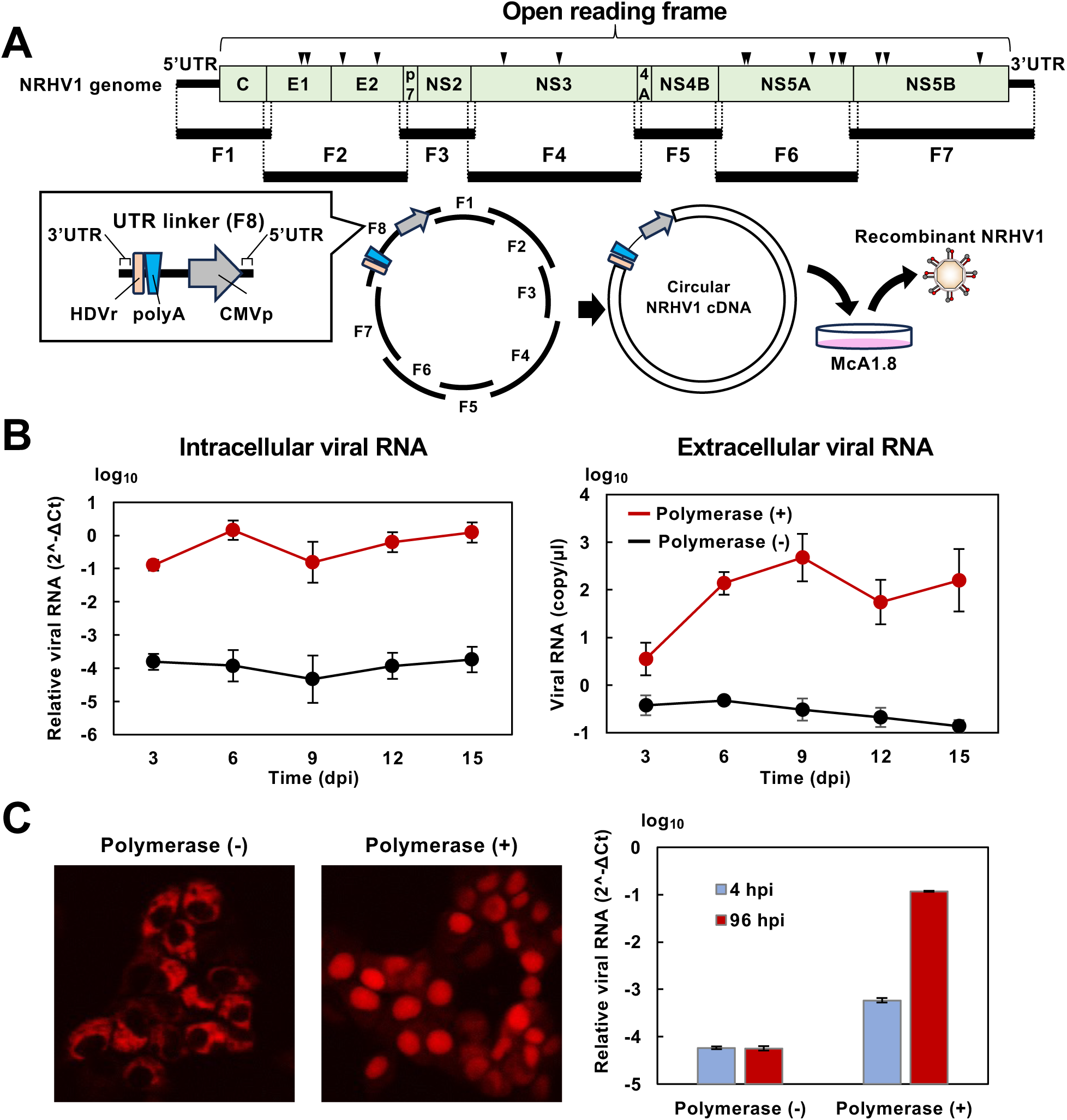
Generation of recombinant NRHV1 by CPER in cultured rat hepatoma-derived cells. (A) Schematic representation of the generation of a recombinant NRHV1 genome by CPER. The full-length NRHV1 genome was divided into seven partially overlapping fragments (F1 to F7). These fragments were subjected to CPER to generate a circularized NRHV1 genome. Circularized DNA was transfected into McA1.8 cells to recover virions. Arrowheads indicate the genomic positions of the cell-adaptive mutations listed in Table 1. HDVr: hepatitis delta virus ribozyme; polyA: polyadenylation signal; CMVp: cytomegalovirus promoter. (B) Time course of intracellular (left) and extracellular (right) viral RNA levels in McA1.8 cells transfected with circularized DNA generated by CPER (polymerase +). Cells transfected with the DNA preparation subjected to CPER without DNA polymerase were used as a negative control (polymerase –). (C) Culture supernatants from the cells transfected with the indicated CPER preparations were collected at 15 dpt. A 0.1-ml aliquot of each supernatant was inoculated into the indicator cell line McA1.8/mCnls-MAVS. The fluorescent mCherry signal was observed at 96 hpi (left). Intracellular NRHV1 RNA levels were quantified at 4 hpi and 96 hpi (right).

**Table 1.** List of cell culture–adaptive mutations.

| Protein | Pos | Ref | Alt | AA |
| --- | --- | --- | --- | --- |
| E1 | 1386 | A | G | M301V |
|  | 1455 | A | G | T324A |
| E2 | 1849 | T | C | L455P |
|  | 2242 | T | C | L586S |
| NS3 | 3664 | A | G | K1060R |
|  | 4287 | A | G | T1268A |
| NS5A | 6375 | G | A | V1964I |
|  | 6408 | G | A | A1975T |
|  | 7135 | T | C | F2217S |
|  | 7362 | T | C | S2293P |
|  | 7459 | A | G | Q2325R |
|  | 7486 | A | G | K2334R |
| NS5B | 7884 | A | G | T2467A |
|  | 7971 | T | C | Y2496N |
|  | 9022 | C | T | A2846V |

The cell-adapted NRHV1 strain previously reported [17] was further passaged seven times in McA1.8 cells, and the resulting strain was designated as rn-1cp7. Mutations identified by comparing the genomic sequence of rn-1cp7 with that of the prototype rn-1 are summarized in the table. “Pos,” “Ref,” and “Alt” indicate the nucleotide position, the nucleotide of the prototype, and the nucleotide of rn-1cp7, respectively. “AA” indicates the corresponding amino acid substitution. For example, a substitution from methionine to valine at amino acid residue 301 is denoted as M301V.

Next, we compared recombinant NRHV1 (rNRHV1) with parental NRHV1 (pNRHV1; strain rn-1cp7) in terms of the localization of double-stranded RNA (dsRNA) and NS5A, the expression of NS5A and the core protein, and sensitivity to direct-acting antivirals (DAAs). The supernatant from cells transfected with the CPER product was inoculated into naïve McA1.8 cells, and the viral inoculation was passaged four times to amplify viral titers (Supplementary Figure 1). In rNRHV1-infected cells, NS5A partially colocalized with dsRNA, similar to the localization pattern observed in pNRHV1-infected cells. NS5A protein appeared as granular structures in the cytoplasm of both rNRHV1- and pNRHV1-infected cells (Figure 2A). The expression levels and molecular sizes of NS5A and the core protein, as well as the phosphorylation pattern of NS5A, in rNRHV1-infected cells were comparable to those in pNRHV1-infected cells (Figure 2B). The phosphorylation pattern of NS5A and the sensitivities to pibrentasvir (PIB) and daclatasvir (DCV) were also similar between rNRHV1 and pNRHV1 (Figure 2 C, D). The IC₅₀ values of PIB for rNRHV1 and pNRHV1 were 1.12 µM and 1.64 µM, respectively, while those of DCV were 4.07 µM and 7.33 µM, respectively. These results indicate that recombinant NRHV1 was successfully generated by CPER.

**Figure 2.**
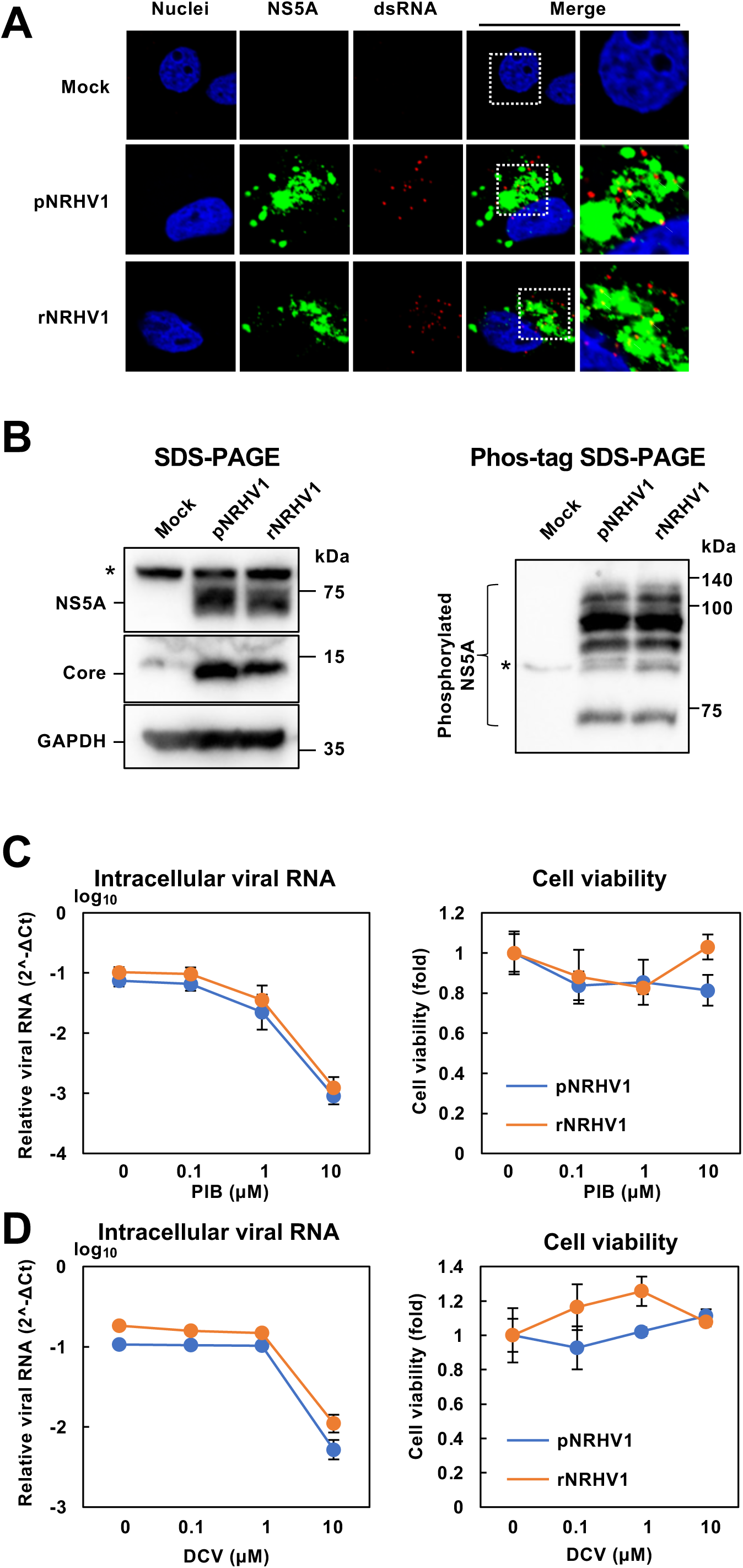
Characterization of parental and recombinant NRHV1. (A) McA1.8 cells were infected with the parental NRHV1 viral stock (pNRHV1), whose mutation profile relative to that of the prototype strain is listed in Table 1, or with recombinant NRHV1 (rNRHV1) at 10 copies/cell. The cells were fixed at 48 hpi and subjected to immunofluorescence analysis. The localization of the NS5A protein and double-stranded RNA (dsRNA) was examined by fluorescence microscopy. Magnified merged images are shown on the right. Arrowheads indicate intracellular regions where NS5A and dsRNA signals colocalize. (B) McA1.8 cells were infected with pNRHV1 or rNRHV1 at 10 copies/cell or mock-treated with medium. At 96 hpi, the cells were analyzed by Western blotting for NS5A and core proteins (left half) and by Mn²⁺-Phos-tag SDS–PAGE (right half). Astarisks indicate nonspecific bands. (C, D) McA1.8 cells were infected with pNRHV1 or rNRHV1 at 10 copies/cell and incubated in the presence of anti-HCV drugs (D: pibrentasvir, PIB; E: daclatasvir, DCV). Intracellular NRHV1 RNA levels were quantified at 96 hpi. Cell viability was evaluated in parallel using an MTS assay.

### Construction of rNRHV1 carrying the HiBiT gene

It has been reported that domain III of HCV NS5A is partially dispensable for replication and can tolerate both partial deletions and insertions of heterologous sequences [18, 19]. Therefore, we next attempted to construct rNRHV1 carrying a reporter gene. The structure of NRHV1 NS5A was predicted using AlphaFold2, and its three domains were provisionally defined on the basis of structural correlations with HCV NS5A. Domain III of NRHV1 NS5A was predicted to be located within amino acid residues 377–506, which is largely disordered and contains a short helical segment, similar to domain III of HCV NS5A (Figure 3A).

**Figure 3.**
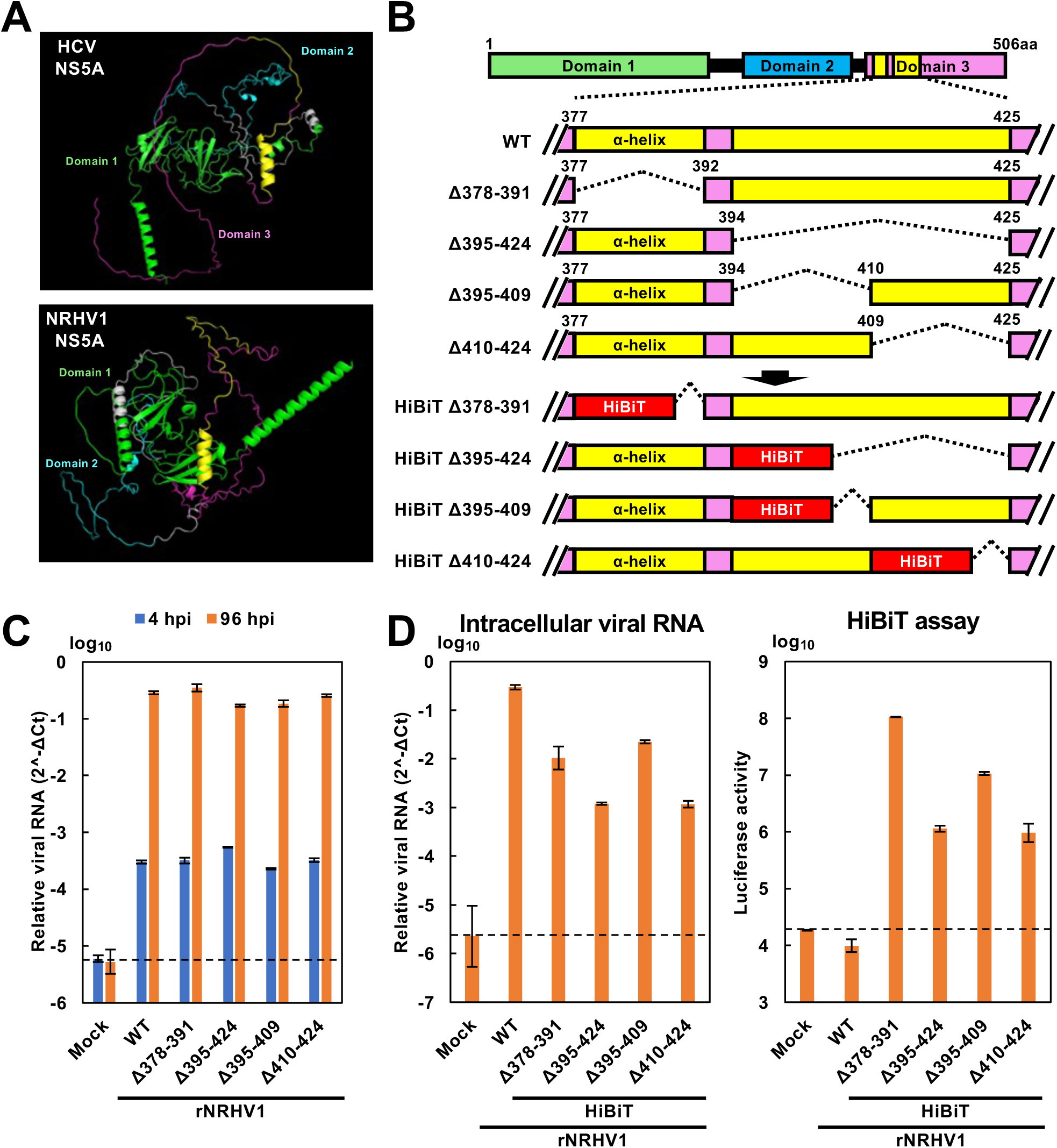
Generation of rNRHV1 carrying either a partial deletion or a HiBiT insertion within the NS5A domain III–coding region. (A) Structural prediction of NS5A proteins by AlphaFold2. NS5A structures of HCV (top) and NRHV1 (bottom) are shown. Green, blue, magenta, and yellow represent domains I, II, III, and the α-helical region spanning residues 378–392, respectively. (B) Schematic representation of deleted regions within the NS5A domain III or insertion sites of the HiBiT sequence. (C) Replication of the deletion mutants. McA1.8 cells were infected with mock, rNRHV1 wild-type (WT), or NS5A domain III deletion mutants at 10 copies/cell. Intracellular NRHV1 RNA levels were quantified at 4 hpi and 96 hpi. (D) Replication of the HiBiT-reporter viruses. McA1.8 cells were infected with mock, rNRHV1 WT, or rNRHV1 carrying a HiBiT sequence inserted at the indicated regions of NS5A domain III (rNRHV1-HiBiT) at 10 copies/cell. Intracellular NRHV1 RNA levels and luciferase activity were measured at 96 hpi. The dotted line indicates the limit of detection (LOD).

Four deletion mutants of rNRHV1 were generated, as shown in Figure 3B. NRHV1 variants with deletions spanning amino acid residues 378–391, 395–424, 395–409, and 410–424 were designated rNRHV Δ378–391, rNRHV Δ395–424, rNRHV Δ395–409, and rNRHV Δ410–424, respectively. These cDNAs were synthesized by CPER and transfected into McA1.8 cells. Supernatants from each mutant were collected at 96 hours post-transfection (hpt) and used to inoculate McA1.8 cells at 10 copies per cell. Intracellular viral RNA was harvested at 4 and 96 hours post-infection (hpi). The viral RNA levels of these deletion variants were comparable to those of wild-type rNRHV1 (rNRHV1 WT) (Figure 3C).

The HiBiT gene was then inserted into the deleted regions of each variant (Figure 3B). The resulting recombinant viruses—based on the rNRHV Δ378–391, rNRHV Δ395–424, rNRHV Δ395–409, and rNRHV Δ410–424 backbones—were designated rNRHV HiBiT Δ378–391, rNRHV HiBiT Δ395–424, rNRHV HiBiT Δ395–409, and rNRHV HiBiT Δ410–424, respectively. These variant cDNAs were generated by CPER and transfected into McA1.8 cells. The culture supernatants were collected at 96 hpt and inoculated into McA1.8 cells at a concentration of 10 copies/cell.

The intracellular RNA level of rNRHV1 WT was higher than that of the other variants; however, no luciferase activity was detected in cells infected with rNRHV1 WT (Figure 3D). Among the HiBiT variants, rNRHV HiBiT Δ378–391 and rNRHV HiBiT Δ395–409 had higher intracellular viral RNA levels and luciferase activities than the other two variants did (Figure 3D). Upon serial passaging, the viral RNA replication of rNRHV HiBiT Δ378–391 and rNRHV HiBiT Δ395–409 slightly increased and plateaued or slightly decreased after the third passage (Figure 4A). Similarly, luciferase activity in cells infected with rNRHV-HiBiT Δ378–391 and rNRHV-HiBiT Δ395–409 was greater than that in cells infected with the other variants (Figure 4A).

**Figure 4.**
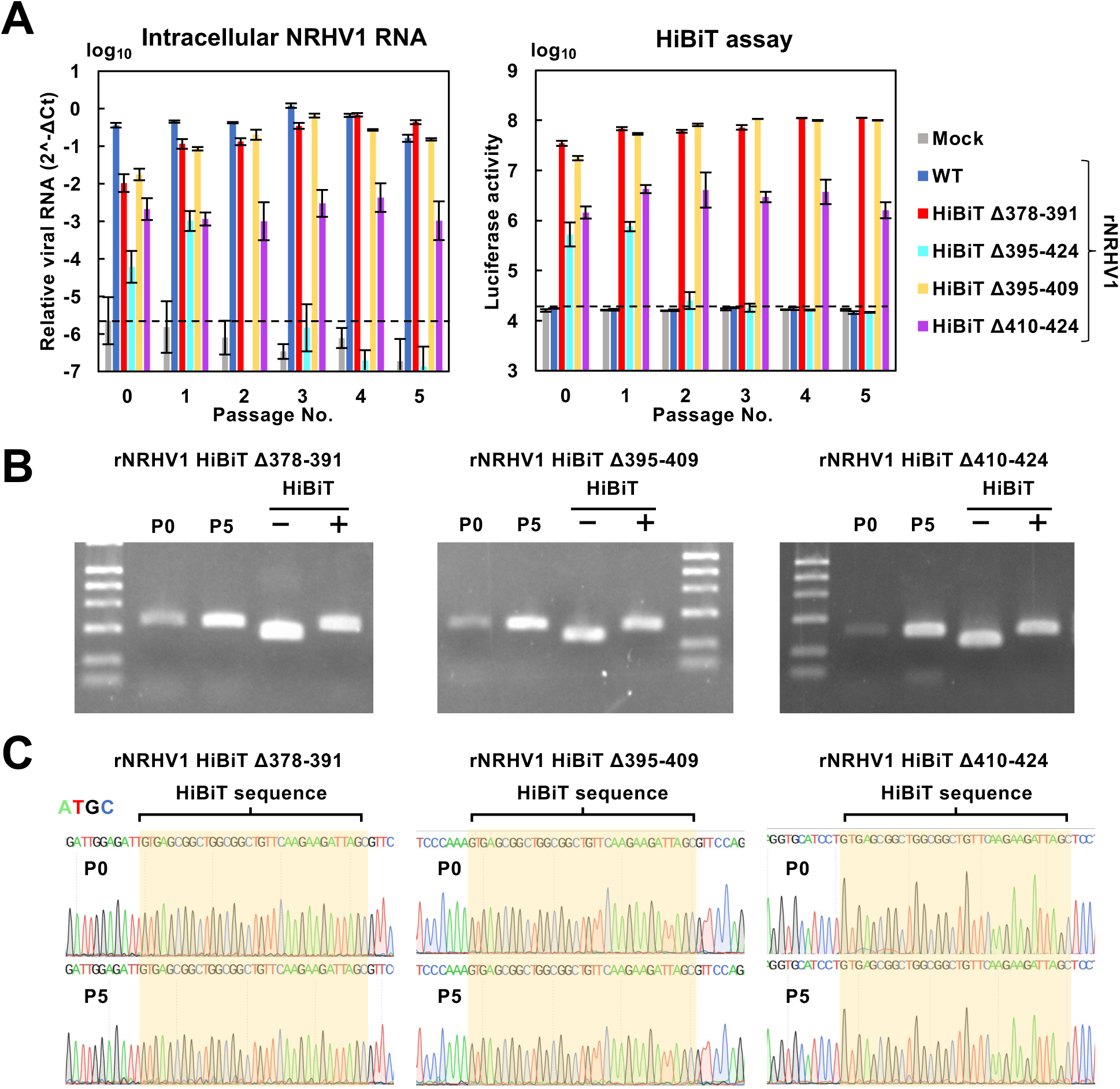
Stability of rNRHV1-HiBiT in vitro. (A) Replication of the HiBiT-reporter viruses. Culture supernatants collected from cells transfected with the indicated CPER products at 6 dpt were designated passage 0 (P0). Naïve McA1.8 cells were inoculated with P0 supernatants to generate P1 supernatants at 96 hpi. This process was repeated sequentially to obtain supernatants from P1 through P5. Naïve McA1.8 cells were then infected with 100 µl of each passage supernatant (P0–P5). Intracellular NRHV1 RNA levels and luciferase activity were quantified at 96 hpi. The dotted line indicates the limit of detection (LOD). (B) Detection of the HiBiT gene. RT–PCR was performed using total RNA extracted from the P0 and P5 supernatants of rNRHV1-HiBiT viruses carrying the HiBiT insertion in the Δ378–391, Δ395–409, or Δ410–424 regions. RT–PCR products were analyzed by agarose gel electrophoresis along with size controls. The size controls consisted of PCR products amplified using F6 fragments with or without the HiBiT insert as templates. (C) Sequence analysis around the inserted HiBiT region. PCR products obtained from RT–PCR in (B) for P0 and P5 samples were subjected to sequencing.

The HiBiT gene was retained in the genomes of rNRHV HiBiT Δ378–391, rNRHV HiBiT Δ395–409, and rNRHV HiBiT Δ410–424 (Figure 4B and C). Replacing the HiBiT gene with regions spanning residues 378–391 or 395–409 did not substantially impair viral propagation compared with that of the other mutants (Figure 4B and C). Although we also attempted to generate rNRHV1 variants carrying the NanoLuc gene instead of the HiBiT gene, all of these variants resulted in extremely low viral RNA levels and a passage-dependent reduction in luciferase activity (Supplementary Figure 2). Thus, rNRHV1 HiBiT Δ378–391 was selected for subsequent experiments because of its stability and replication efficiency.

### Characterization of rNRHV1 HiBiT Δ378–391

We examined the effect of HiBiT insertion on viral growth in a cell culture model. As shown in Supplementary Figure 1, the culture supernatant from cells transfected with CPER-derived rNRHV1 HiBiT Δ378–391 was inoculated into naïve McA1.8 cells, followed by four serial passages to amplify the viral titer as a stock, similar to the procedure used for rNRHV1 WT. Naïve McA1.8 cells were then inoculated with either rNRHV1 WT or rNRHV1 HiBiT Δ378–391 at 10 copies/cell. Intracellular viral RNA levels and luciferase activity were measured at 4, 24, 48, 72, and 96 hours post-infection (hpi) (Figure 5A). Compared with that of rNRHV1 WT, the intracellular RNA level of rNRHV1 HiBiT Δ378–391 increased more slowly but reached a comparable level at 96 hpi (Figure 5A, left graph). The luciferase activity of cells infected with rNRHV1 HiBiT Δ378–391 increased in a time-dependent manner corresponding to the viral RNA levels, whereas no luciferase activity was detected in cells infected with rNRHV1 WT (Figure 5A, right graph). We next examined the intracellular localization of NS5A in cells infected with either rNRHV1WT or rNRHV1 HiBiT Δ378–391. In cells infected with rNRHV1 HiBiT Δ378–391, NS5A displayed a granular distribution pattern similar to that observed in rNRHV1 WT-infected cells (Figure 5B). The fluorescence signals detected with anti-NS5A and anti-HiBiT antibodies overlapped in cells infected with the mutant virus (Figure 5B). NS5A expression levels in cells infected with rNRHV1 HiBiT Δ378–391 were comparable to those in rNRHV1 WT–infected cells, whereas HiBiT expression was detected only in rNRHV1 HiBiT Δ378–391-infected cells (Figure 5C). The lower molecular weight of HiBiT-inserted NS5A than that of native NS5A (Figure 5C, left half) may be attributable to differences in phosphorylation levels (Figure 5C, right half). Taken together, these data suggest that the insertion of HiBiT into NS5A slightly delays viral growth.

**Figure 5.**
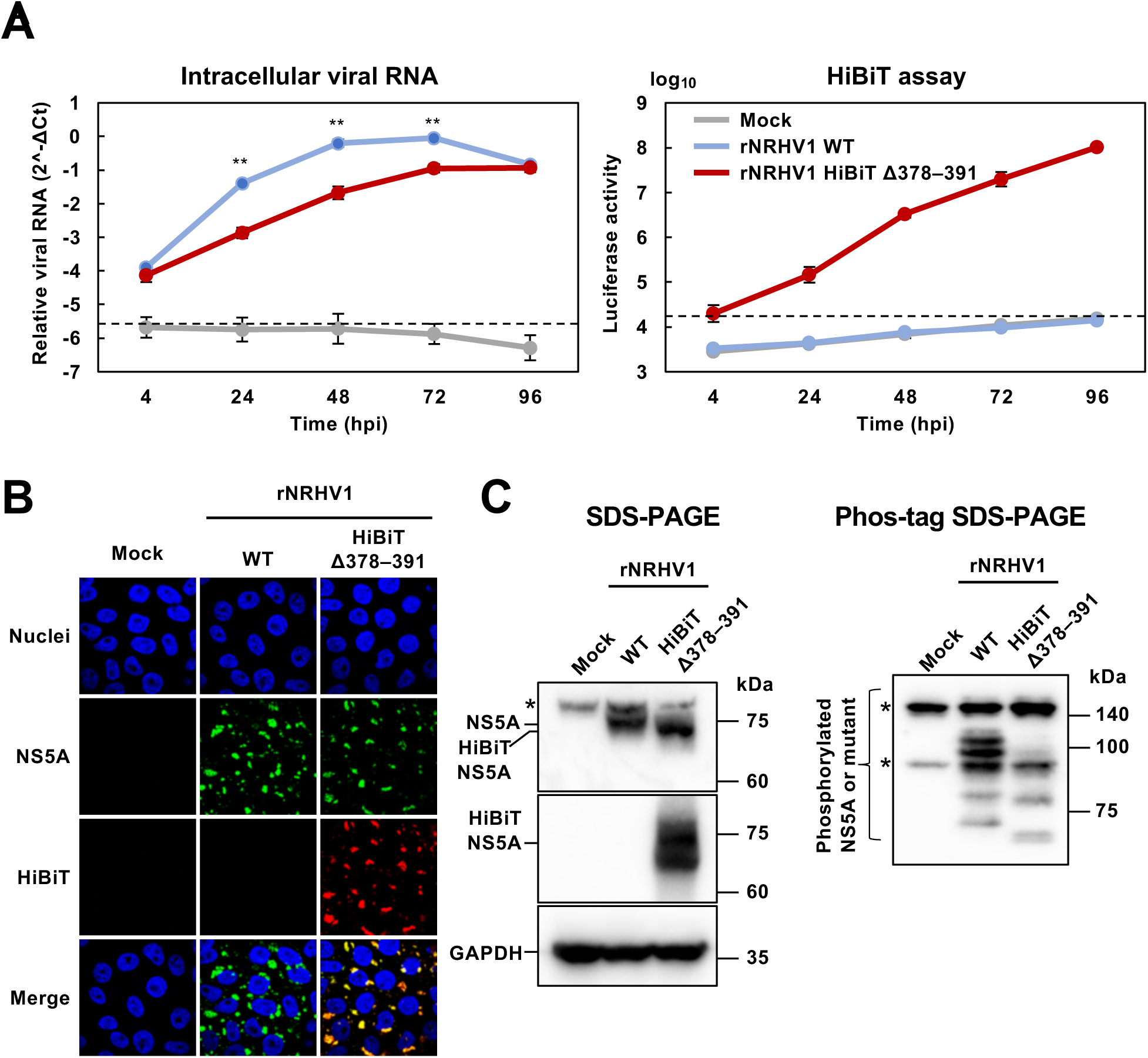
Effect of HiBiT insertion on viral characteristics. McA1.8 cells were infected with rNRHV1 WT or rNRHV1-HiBiT Δ378–391 at 10 copies/cell. (A) Time course of viral replication and luciferase activity. Intracellular NRHV1 RNA levels and luciferase activity were measured at the indicated time points. Asterisks indicate statistically significant differences between the wild-type and mutant viruses. **: p < 0.01. (B) Localization of NS5A in infected cells. The localization of the NS5A and HiBiT epitopes was analyzed by immunofluorescence staining. (C) Western blot analysis of lysates extracted from cells infected with the indicated viruses or mock control at 96 hpi. (left half) The same lysates were subjected to Mn²⁺-Phos-tag SDS–PAGE to assess NS5A phosphorylation (right half). Asterisks indicate nonspecific bands. HiBiT NS5A indicates that the NS5A 378–391 region was replaced with HiBiT.

### Application of rNRHV1 HiBiT Δ378–391 for Evaluating Antiviral Drug and Neutralizing Antibody Activity

The NanoBiT system can be useful for assessing antiviral drug activity and neutralizing antibody activity [20, 21]. We investigated whether rNRHV1 HiBiT Δ378–391 could be used to monitor the efficacy of antiviral agents using luciferase activity as an indicator of viral RNA levels. When naïve McA1.8 cells were inoculated with either rNRHV1 WT or rNRHV1 HiBiT Δ378–391, treatment with PIB or DCV reduced luciferase activity in rNRHV1 HiBiT Δ378–391-infected cells, which was consistent with the decrease in viral RNA levels (Figure 6). The intracellular RNA levels decreased upon treatment with PIB or DCV (Figure 6). The IC₅₀ values of PIB were 1.12 µM for rNRHV1 WT and 0.13 µM for rNRHV1 HiBiT Δ378–391, whereas those of DCV were 3.71 µM and 0.35 µM, respectively. These results suggest that the antiviral effects of DAAs are more pronounced against rNRHV1 HiBiT Δ378–391 than against rNRHV1 WT. We next examined whether the HiBiT system could also be applied to evaluate neutralizing antibody activity. Serum was obtained from SD rats infected with NRHV1 at 26 weeks post-infection (wpi) or from naïve SD rats and then heat inactivated. These serum samples were incubated with rNRHV1 HiBiT Δ378–391 for 1 h at 37°C and subsequently inoculated into naïve McA1.8 cells at 5 copies/cell. Luciferase activity was measured at 48 hpi. Preincubation of the virus with 100-fold–diluted serum before infection reduced luciferase activity to less than 40% of that observed with mock serum (Figure 7A and B). The antiviral effect of serum from infected rats was dependent on serum concentration (Figure 7B). In contrast, the addition of serum from infected rats to the culture supernatant at 4 hpi did not affect luciferase activity (Figure 7B). The serum from infected rats contained antibodies against both envelope and nonstructural proteins (Figure 7C). Together, these findings indicate that our HiBiT-based system is applicable for antiviral assays targeting both small-molecule compounds and neutralizing antibodies.

**Figure 6.**
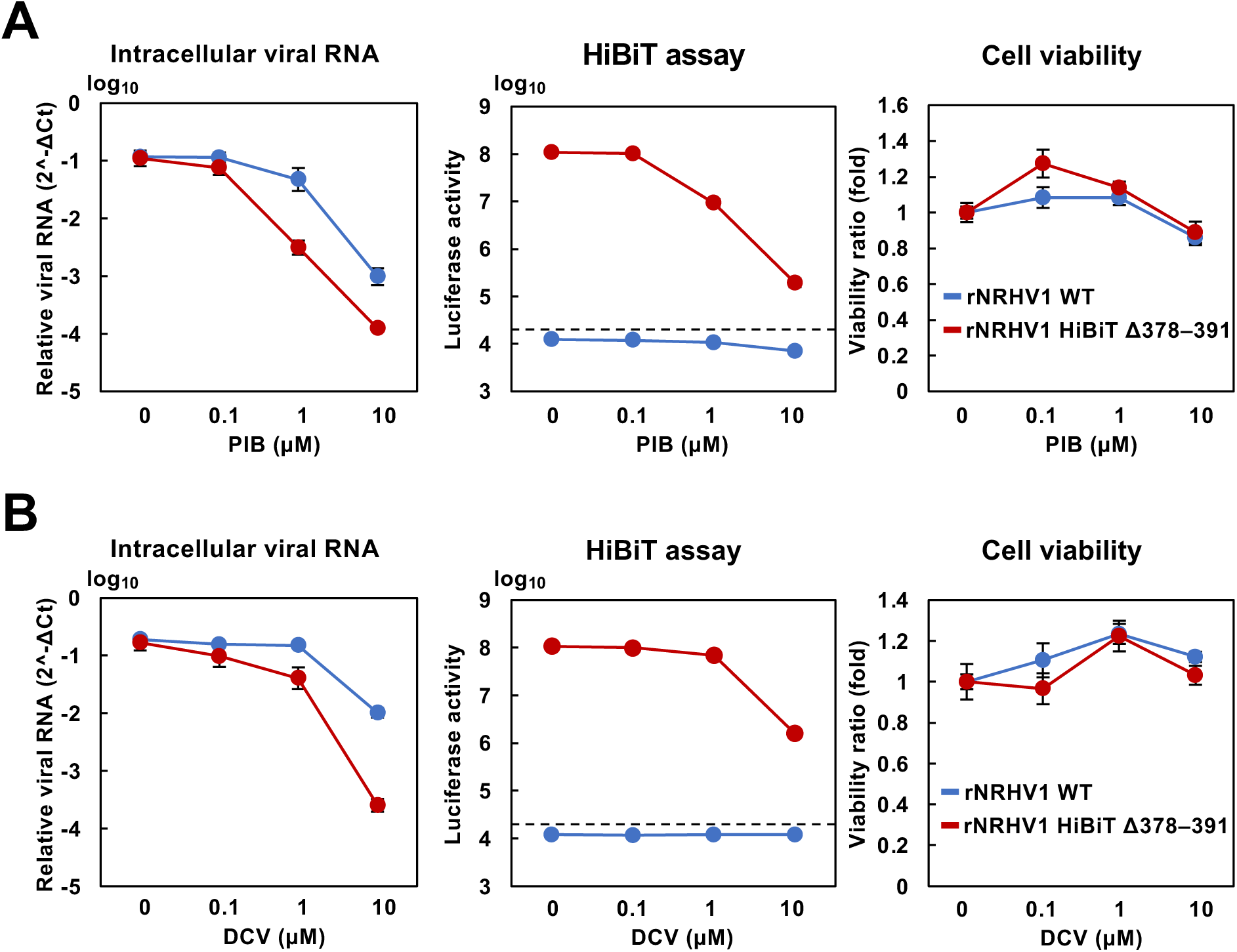
Antiviral sensitivities of rNRHV1 WT and rNRHV1-HiBiT Δ378–391. McA1.8 cells were infected with rNRHV1 WT or rNRHV1-HiBiT Δ378–391 at 10 copies/cell and incubated in the presence of PIB (A) or DCV (B). Intracellular NRHV1 RNA levels, luciferase activities, and cell viability were measured at 96 hpi, as described in the Materials and Methods. The dotted line indicates the limit of detection (LOD).

**Figure 7.**
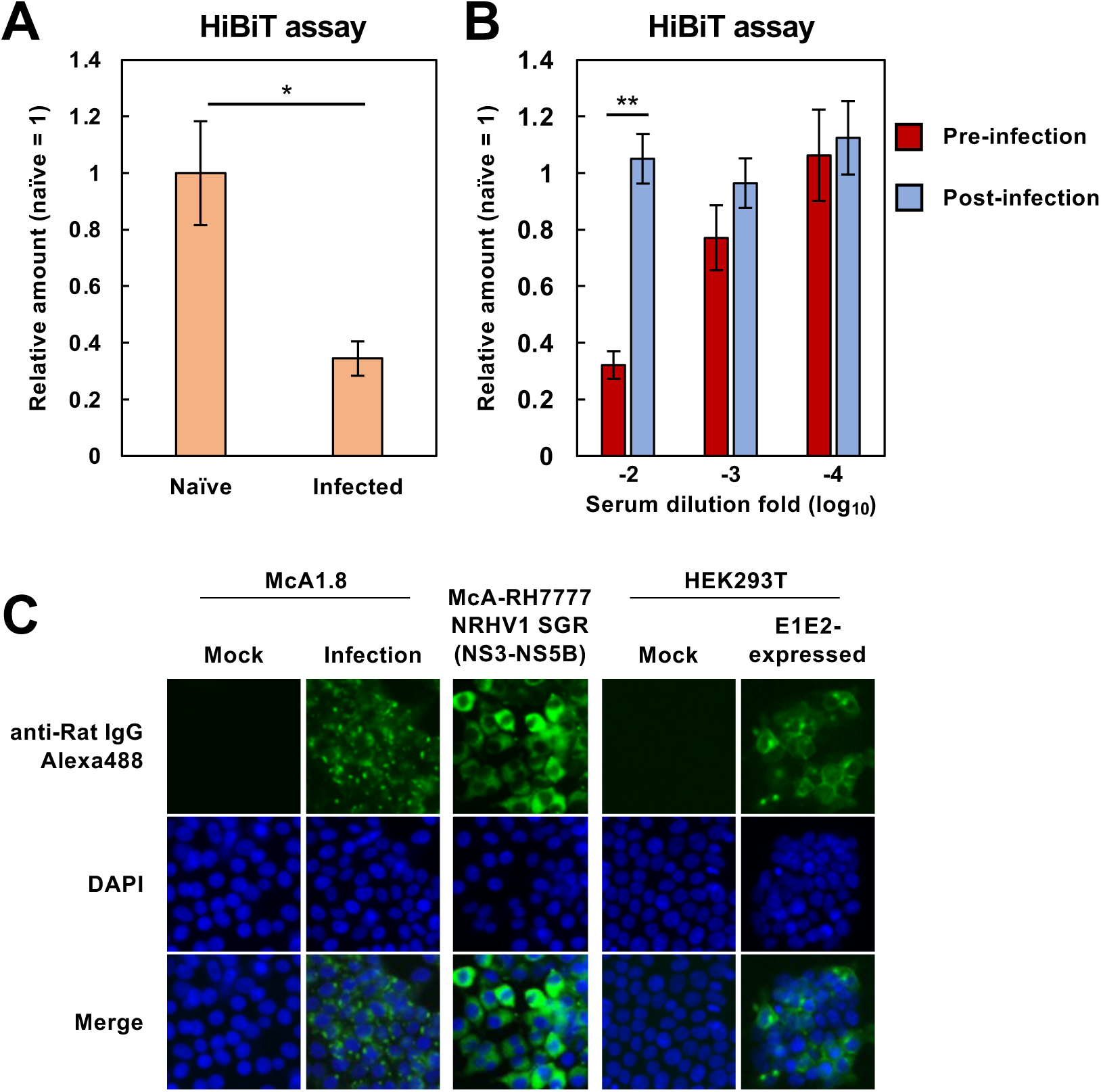
Effects of rat serum on viral growth. (A) rNRHV1-HiBiT Δ378–391 was incubated for 1 h at 37°C with serum obtained from NRHV1-infected SD rats at 26 wpi or from naïve rats. McA1.8 cells were then infected with the virus–serum mixtures at 5 copies/cell. Luciferase activity was measured at 48 hpi. *: p < 0.05. (B) Effect of serum postincubation on viral growth. Serum diluted to the indicated folds was either incubated with the virus preparation prior to infection (preinfection) or added to cells at 4 hpi (postinfection). Luciferase activity was quantified at 48 hpi. Relative HiBiT activities were normalized to the corresponding naïve-serum controls for the preinfection and postinfection groups. **: p < 0.01. (C) Reactivity of the antiserum to viral proteins. Serum obtained from NRHV1-infected SD rats at 26 wpi was used for immunofluorescence assays with HEK293T cells expressing NRHV1 E1-E2, subgenomic replicon-harboring McA1.8 cells, and NRHV1-infected McA1.8 cells.

### rNRHV1 HiBiT Δ378–391 causes persistent infection in vivo

Next, we examined the in vivo infectivity and stability of rNRHV1 HiBiT Δ378–391. Recombinant viruses were inoculated into NOD–SCID mice. In mice infected with WT rNRHV1, serum viral RNA levels remained at approximately 10⁴–10⁵ copies/µl of blood from 4 to 8 wpi. In contrast, mice infected with rNRHV1 HiBiT Δ378–391 presented variable serum viral RNA levels between 2 and 6 wpi, which stabilized at 8 wpi (Figure 8A). Liver samples were collected from mice infected with each virus at 8 wpi. When viral RNA levels and luciferase activities were measured in the livers of the individual groups, both signals were detected in mice infected with rNRHV1 HiBiT Δ378–391 (Figure 8B). In contrast, viral RNA was detected but luciferase activity was absent in the livers of the rNRHV1 WT–infected mice (Figure 8B). Luciferase activity was detected exclusively in the liver and not in the intestine, spleen, lung, testis, salivary gland, kidney, or brain (Supplementary Figure 3), suggesting that rNRHV1 HiBiT Δ378–391 retains liver tropism similar to that of the prototype NRHV1, as previously reported [11]. Histological analysis revealed viral RNA in liver sections from both rNRHV1 WT– and rNRHV1 HiBiT Δ378–391-infected mice. However, anti-HiBiT antibody-reactive signals were detected in liver tissues from rNRHV1 HiBiT Δ378–391-infected mice but not in those from rNRHV1 WT-infected mice (Figure 8C). Sera obtained from these mouse groups were inoculated into naïve McA1.8 cells. Viral RNA was detected in all inoculated cells, whereas luciferase activity was observed only in cells inoculated with serum from rNRHV1 HiBiT Δ378–391-infected mice (Figure 8D). However, luciferase activity was relatively low in cells infected with serum from two of the five rNRHV1 HiBiT Δ378–391-infected mice. To determine the NRHV1 genome sequences present in the culture supernatants and inoculated sera, total RNA was extracted from both. Sera from mice No. 1, 2, and 3 infected with rNRHV1 HiBiT Δ378–391 exhibited high infectivity and high luciferase activity in McA1.8 cells, comparable to those from rNRHV1 WT-infected mice (Figure 8D). In contrast, sera from mice No. 4 and 5 showed high viral RNA levels but very low luciferase activity in McA1.8 cells (Figure 8D). An analysis of the HiBiT gene retention rates in the viral RNA genomes within these serum samples revealed values of 95%, 96%, 94%, 5%, and 0.3% for mice No. 1, 2, 3, 4, and 5, respectively. Furthermore, several additional mutations were identified in serum-derived rNRHV1 WT and rNRHV1 HiBiT Δ378–391 viruses that were not detected in the corresponding viruses passaged in cell culture, although these mutations were not uniformly present across all mice (Table 2). These results suggest that rNRHV1 HiBiT Δ378–391 is capable of infecting both cultured cells and mice and can maintain the HiBiT gene but may occasionally lose it in vivo.

**Figure 8.**
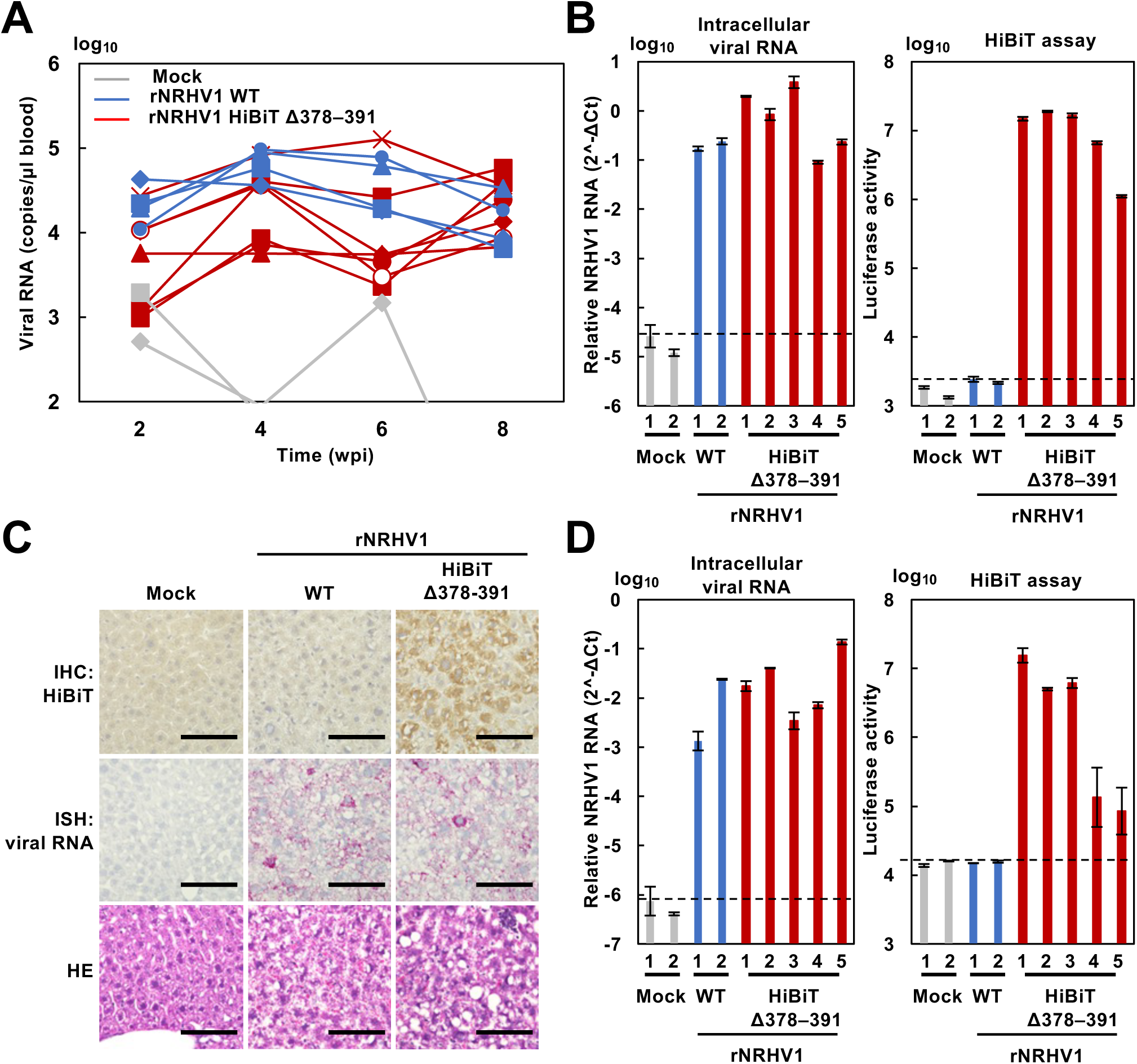
Growth efficiency of rNRHV1 in NOD–SCID mice. NOD–SCID mice were intravenously injected with 8 × 10⁵ copies of rNRHV1 WT (n = 4), rNRHV1-HiBiT Δ378–391 (n = 7), or medium (mock, n = 3). Viral replication was monitored up to 8 wpi. (A) Time course of viral RNA levels in blood, as quantified by qRT–PCR. (B) Viral RNA levels and luciferase activity in liver tissues collected at 8 wpi. (C) Detection of the NS5A antigen and NRHV1 RNA in liver tissues collected at 8 wpi. NS5A and viral RNA were detected by immunohistochemistry using an anti-HiBiT antibody (top panels) and by in situ hybridization using probes targeting the NS3 region (middle panels), respectively. Histopathological examination was performed using H&E-stained sections (bottom panels). H&E: hematoxylin and eosin. Scale bars: 100 µm. (D) Infectivity of mouse-derived viruses in cell culture. McA1.8 cells were inoculated with serum collected at 8 wpi, and intracellular NRHV1 RNA levels and luciferase activity were measured at 96 hpi. The dotted line indicates the limit of detection (LOD).

**Table 2.**
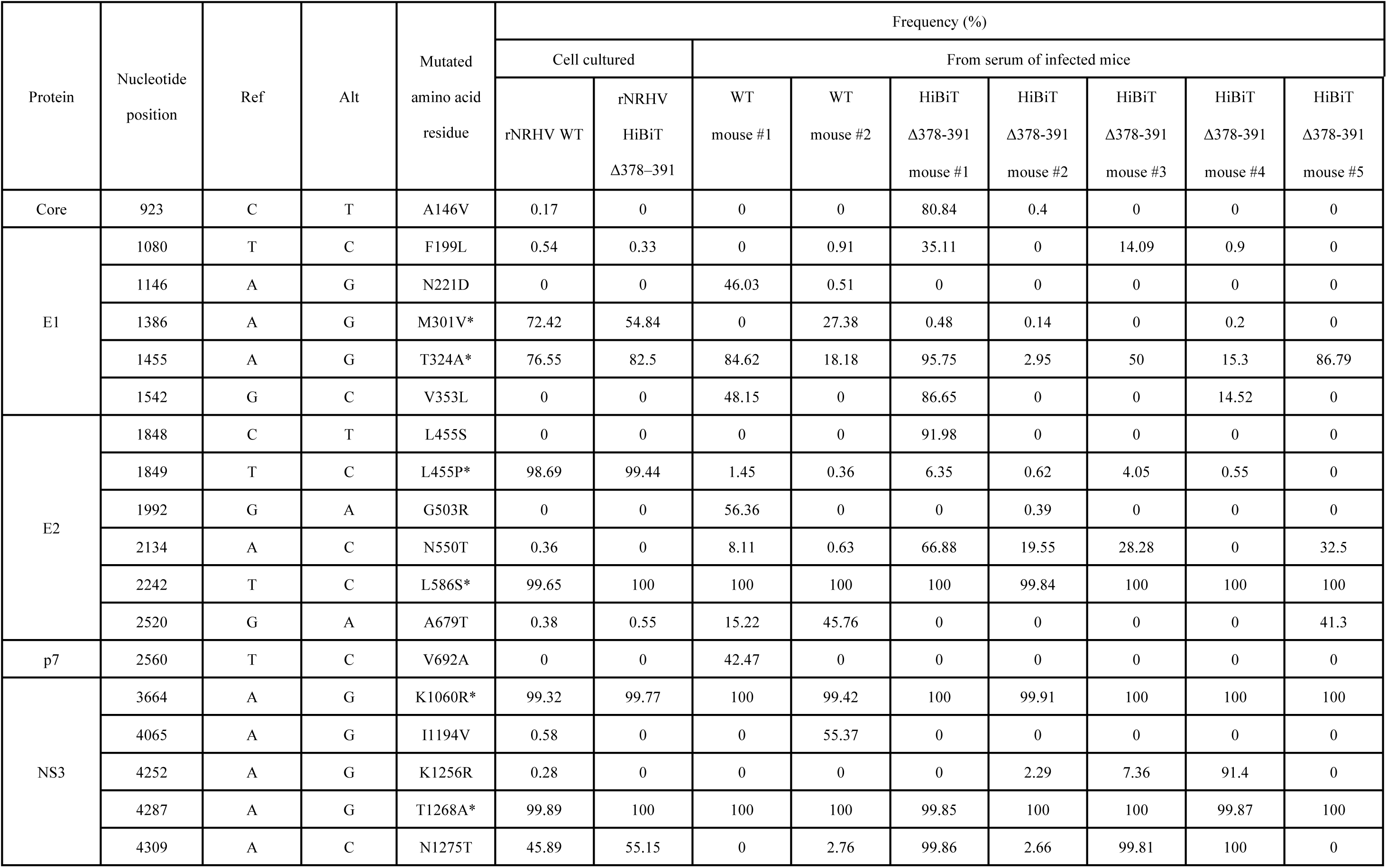
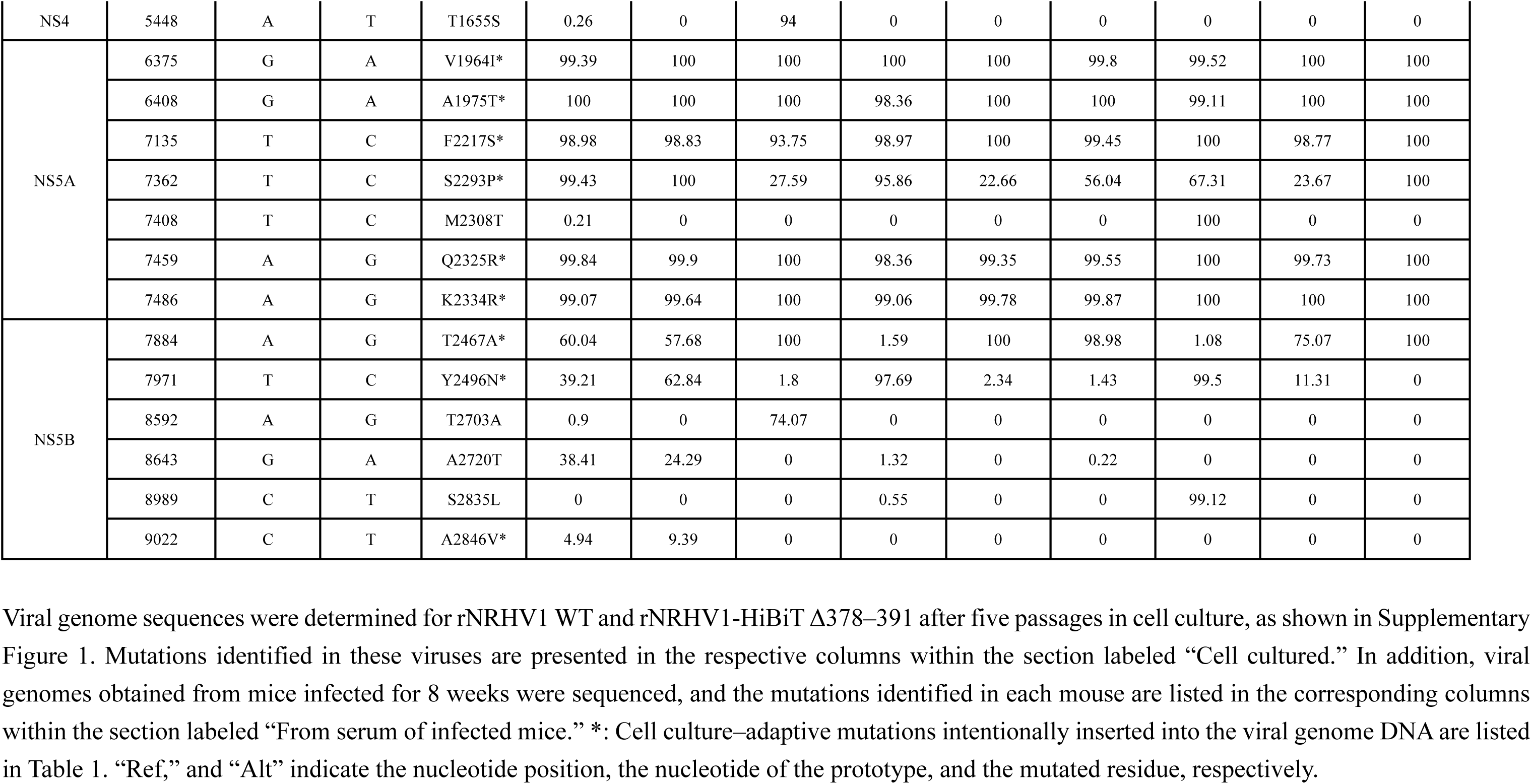
List of viral mutations identified in viral RNAs derived from cell culture and from sera of rNRHV1-infected mice.

| Protein | Nucleotide position | Ref | Alt | Mutated amino acid residue | Frequency (%) |  |  |  |  |  |  |  |  |
| --- | --- | --- | --- | --- | --- | --- | --- | --- | --- | --- | --- | --- | --- |
|  |  |  |  |  | Cell cultured |  | From serum of infected mice |  |  |  |  |  |  |
|  |  |  |  |  | rNRHV WT | rNRHV HiBiT<br>Δ378–391 | WT mouse #1 | WT mouse #2 | HiBiT Δ378-391 mouse #1 | HiBiT Δ378-391 mouse #2 | HiBiT Δ378-391 mouse #3 | HiBiT Δ378-391 mouse #4 | HiBiT Δ378-391 mouse #5 |
| Core | 923 | C | T | A146V | 0.17 | 0 | 0 | 0 | 80.84 | 0.4 | 0 | 0 | 0 |
| E1 | 1080 | T | C | F199L | 0.54 | 0.33 | 0 | 0.91 | 35.11 | 0 | 14.09 | 0.9 | 0 |
|  | 1146 | A | G | N221D | 0 | 0 | 46.03 | 0.51 | 0 | 0 | 0 | 0 | 0 |
|  | 1386 | A | G | M301V* | 72.42 | 54.84 | 0 | 27.38 | 0.48 | 0.14 | 0 | 0.2 | 0 |
|  | 1455 | A | G | T324A* | 76.55 | 82.5 | 84.62 | 18.18 | 95.75 | 2.95 | 50 | 15.3 | 86.79 |
|  | 1542 | G | C | V353L | 0 | 0 | 48.15 | 0 | 86.65 | 0 | 0 | 14.52 | 0 |
| E2 | 1848 | C | T | L455S | 0 | 0 | 0 | 0 | 91.98 | 0 | 0 | 0 | 0 |
|  | 1849 | T | C | L455P* | 98.69 | 99.44 | 1.45 | 0.36 | 6.35 | 0.62 | 4.05 | 0.55 | 0 |
|  | 1992 | G | A | G503R | 0 | 0 | 56.36 | 0 | 0 | 0.39 | 0 | 0 | 0 |
|  | 2134 | A | C | N550T | 0.36 | 0 | 8.11 | 0.63 | 66.88 | 19.55 | 28.28 | 0 | 32.5 |
|  | 2242 | T | C | L586S* | 99.65 | 100 | 100 | 100 | 100 | 99.84 | 100 | 100 | 100 |
|  | 2520 | G | A | A679T | 0.38 | 0.55 | 15.22 | 45.76 | 0 | 0 | 0 | 0 | 41.3 |
| p7 | 2560 | T | C | V692A | 0 | 0 | 42.47 | 0 | 0 | 0 | 0 | 0 | 0 |
| NS3 | 3664 | A | G | K1060R* | 99.32 | 99.77 | 100 | 99.42 | 100 | 99.91 | 100 | 100 | 100 |
|  | 4065 | A | G | I1194V | 0.58 | 0 | 0 | 55.37 | 0 | 0 | 0 | 0 | 0 |
|  | 4252 | A | G | K1256R | 0.28 | 0 | 0 | 0 | 0 | 2.29 | 7.36 | 91.4 | 0 |
|  | 4287 | A | G | T1268A* | 99.89 | 100 | 100 | 100 | 99.85 | 100 | 100 | 99.87 | 100 |
|  | 4309 | A | C | N1275T | 45.89 | 55.15 | 0 | 2.76 | 99.86 | 2.66 | 99.81 | 100 | 0 |
| NS4 | 5448 | A | T | T1655S | 0.26 | 0 | 94 | 0 | 0 | 0 | 0 | 0 | 0 |
| NS5A | 6375 | G | A | V1964I* | 99.39 | 100 | 100 | 100 | 100 | 99.8 | 99.52 | 100 | 100 |
|  | 6408 | G | A | A1975T* | 100 | 100 | 100 | 98.36 | 100 | 100 | 99.11 | 100 | 100 |
|  | 7135 | T | C | F2217S* | 98.98 | 98.83 | 93.75 | 98.97 | 100 | 99.45 | 100 | 98.77 | 100 |
|  | 7362 | T | C | S2293P* | 99.43 | 100 | 27.59 | 95.86 | 22.66 | 56.04 | 67.31 | 23.67 | 100 |
|  | 7408 | T | C | M2308T | 0.21 | 0 | 0 | 0 | 0 | 0 | 100 | 0 | 0 |
|  | 7459 | A | G | Q2325R* | 99.84 | 99.9 | 100 | 98.36 | 99.35 | 99.55 | 100 | 99.73 | 100 |
|  | 7486 | A | G | K2334R* | 99.07 | 99.64 | 100 | 99.06 | 99.78 | 99.87 | 100 | 100 | 100 |
| NS5B | 7884 | A | G | T2467A* | 60.04 | 57.68 | 100 | 1.59 | 100 | 98.98 | 1.08 | 75.07 | 100 |
|  | 7971 | T | C | Y2496N* | 39.21 | 62.84 | 1.8 | 97.69 | 2.34 | 1.43 | 99.5 | 11.31 | 0 |
|  | 8592 | A | G | T2703A | 0.9 | 0 | 74.07 | 0 | 0 | 0 | 0 | 0 | 0 |
|  | 8643 | G | A | A2720T | 38.41 | 24.29 | 0 | 1.32 | 0 | 0.22 | 0 | 0 | 0 |
|  | 8989 | C | T | S2835L | 0 | 0 | 0 | 0.55 | 0 | 0 | 99.12 | 0 | 0 |
|  | 9022 | C | T | A2846V* | 4.94 | 9.39 | 0 | 0 | 0 | 0 | 0 | 0 | 0 |

## Discussion

In this study, we established an NRHV1 reverse genetics system based on CPER and successfully generated recombinant viruses (Figure 1). We previously reported an NRHV1-susceptible cell line, McA1.8, and a cell-adapted NRHV1 strain [17]. The cell-adapted viral strain was passaged seven times in McA1.8 cells and designated strain rn-1cp7. The genome sequence was determined and compared with that of the prototype rn-1 strain (KX9015133). Compared with the previously reported adaptive viral genome, five mutations (L455P, L586S, F2217S, Q2325R, K2334R, and A2846V) [17] were conserved after seven passages, although other mutations reverted to the original amino acid residues [17]. L586S, located in the E2-coding region, has also been reported in another cell-adapted viral strain [22], but the others have not.

The virus stock generated by CPER was passaged five times (Supplementary Figure 1) and then used as rNRHV1 WT for subsequent experiments. Additional mutations arising during the five passages are listed in Table 2. To clarify the virological significance of the cell-adaptive mutations identified in this study, we examined their potential functional roles on the basis of structural predictions. L586A and K1060R, located in E2 and NS3, respectively, were found in the cell-adapted strain. These mutations were consistently detected at frequencies greater than 99% in viral genomes from both the rNRHV1 WT and rNRHV1 HiBiT Δ378–391 virus stocks, indicating high genetic stability under both in vitro and in vivo conditions (Table 2).

The substitution of Leu586 with Ser (L586S) was previously reported as a cell culture–adaptive mutation during serial passaging [22]. Although limited information has been reported concerning the structural homology between the NRHV1 and HCV E2 proteins [23, 24], we predicted the NRHV1 E2 structure around Leu586 using AlphaFold2. Leu586 of NRHV1 is likely positioned on an antiparallel β-hairpin within the back layer of E2 and corresponds to Leu644 of HCV (Supplementary Figure 4). Analyses of an E2 mutant derived from the serially passaged HCV JFH1 strain suggest that HCV Leu644 contributes to binding with the scavenger receptor class B type I (SR-BI) [25]. Thus, the mutation of Leu586 to Ser in NRHV1 E2 may influence E2–SR-BI interactions under both in vitro and in vivo conditions.

The structure of NRHV1 NS3 predicted by AlphaFold2 revealed a domain organization comprising protease and helicase domains similar to those of HCV NS3 (PDB ID: 3O8B) (Supplementary Figure 5). Lys1060 is predicted to reside on a β-sheet within the protease domain. In HCV NS3, the protease domain consists of two β-barrel subdomains stabilized by interactions with both Zn²⁺ and the NS4A cofactor [26, 27]. Replacement of Lys1060 with Arg may influence the stability and enzymatic activity of the protease domain.

We next attempted to insert an exogenous gene into the NRHV1 genome and focused on the NS5A-coding region, since the EGFP and NanoLuc genes have been stably inserted into domain III of the HCV NS5A-coding region during viral propagation [28–30], albeit with a slight reduction in viral titer compared with that of the parental strain. To identify regions that are resistant to deletion without severely affecting viral propagation, several deletion mutants were constructed (Figure 3A–C). Deletions corresponding to residues 378–391 (Δ378–391), 395–409 (Δ395–409), 395–424 (Δ395–424), or 410–424 (Δ410–424) of NS5A did not affect viral replication (Figure 3C). However, the insertion of the HiBiT gene into regions encoding residues 378–391 or 395–409 slightly reduced viral propagation, whereas insertion into the other two regions markedly impaired viral propagation (Figure 3D).

The rNRHV1 HiBiT Δ378–391 virus persistently infected three immunodeficient mice, maintaining the HiBiT gene up to 8 wpi, although two mice infected with rNRHV1 lost the HiBiT gene from the replicating virus. Insertion of the NanoLuc gene into the regions encoding residues 378–391, 395–409, 395–424, or 410–424 was unsuccessful (Supplementary Figure 2). Although recombinant HCVs containing the NanoLuc or EGFP genes have been successfully generated, stable recombinant NRHV1 containing NanoLuc or EGFP could not be obtained. This failure is unlikely due to limitations in viral packaging capacity, as both viral RNA and luciferase activity were detected in cells inoculated with supernatants from transfected cells containing the NanoLuc gene (Supplementary Figure 2). Instead, the insertion of an exogenous gene may interfere with the function of NS5A in viral replication or particle formation.

The N-terminal domain I of NS5A, which includes a zinc-binding motif, is conserved among Hepaciviruses, whereas domains II and III exhibit low sequence similarity [31, 32]. Several reporter HCVs have been generated by inserting exogenous genes such as NanoLuc into domain III of NS5A, but in this study, NRHV1-based recombinant viruses carrying genes of a size similar to or larger than NanoLuc could not be constructed. One possible explanation is that structural differences between NRHV1 and HCV in domains II and III affect tolerance for exogenous gene insertion. Reporter HCVs are typically constructed using the genotype 2a strain JFH1 [28–30]. Studies in which the NS5A region of J6/JFH1 was replaced with those from other genotypes have shown decreased replication efficiency and reduced formation of infectious particles [33, 34], suggesting that JFH1-derived NS5A possesses superior functional properties for HCV replication and particle assembly. The insertion of the HiBiT gene was accompanied by a low-frequency substitution (N550T) in the E2-coding region (Table 2), suggesting that HiBiT insertion may influence viral assembly.

In conclusion, we generated a reporter NRHV1 that remains infectious both in vitro and in vivo. Although the size of the inserted reporter gene is limited, this system can be adapted for additional virological applications using other compact reporter constructs. Overall, our system should facilitate future studies on the mechanisms underlying NRHV1 persistence, immunity, and pathogenesis.

## Materials and Methods

### Plasmids

The seven cell-adaptive NRHV1 cDNA (Norway rat hepacivirus 1 rn-1cp7, accession code: LC891941) fragments, covering the entire viral genome and each possessing 40–80 bp overlaps with neighboring fragments, and the UTR linker, containing the 3’-terminal 60 nucleotides of the genome, hepatitis delta virus ribozyme (HDVr), bovine growth hormone polyadenylation signal (BGH polyA), cytomegalovirus promoter (CMVp) and the 5’-terminal 40 nucleotides, were cloned into the pCR-Blunt or pMD20-T vector (Invitrogen, TaKaRa Bio). The HiBiT gene was introduced into the viral genome by PCR using primers containing the HiBiT sequence. The NanoLuc gene was amplified by PCR from the pNL1.1 expression plasmid (Promega).

### Cell lines

All the cell lines were maintained in Dulbecco’s modified Eagle’s medium (DMEM; Sigma–Aldrich) supplemented with 10% fetal bovine serum (FBS; Gibco), nonessential amino acids (Sigma–Aldrich), sodium pyruvate (Sigma–Aldrich), 100 U/ml penicillin (Gibco) and 100 µg/ml streptomycin (Gibco) at 37°C with 5% CO2. The cell line stably expressing mCherry-MAVS was selected with 500 µg/ml zeocin (InvivoGen). The cell line harboring NRHV1 subgenomic replicon (SGR) RNA was maintained in the presence of 500 μg/ml G418 (Nacalai Tesque).

### Generation of the recombinant NRHV1 genome by CPER

Seven DNA fragments (F1 to F7), which were designed for the generation of the entire sequence of NRHV1 (Figure 1A), were synthesized by PCR using the primer pairs described in Table 3. Fragments F1, F2, F3 F4, F5, F6 and F7 were amplified by PCR using the primer pairs NRHV1-F1, NRHV1-F2, NRHV1-F3, NRHV1-F4, NRHV1-F5, NRHV1-F6 and NRHV1-F7, respectively. Each primer had a 40–80 bp overlap sequence at both ends (Figure 1A). The “UTR linker fragment” (F8) consisted of the 3’-terminal UTR (60 nt) of the NRHV1 genome, the hepatitis delta virus ribozyme (HDVr), the polyadenylation signal (polyA), the cytomegalovirus promoter (CMVp) and the 5’-terminal UTR (40 nt) of the NRHV1 genome, in order. Finally, the eight PCR fragments (F1 to F8) were mixed in equal amounts (0.05 pmol each) in 50 μl of the reaction buffer for PrimeSTAR GXL DNA polymerase (Takara Bio, Siga, Japan) and assembled *in vitro* by CPER. CPER was carried out as follows: one cycle of 98°C for 2 min; 20 cycles of 98°C for 10 sec, 55°C for 15 sec and 68°C for 30 min; and one cycle of 68°C for 30 min followed by a hold at 4°C. The resulting products were introduced into McA1.8 cells using Lipofectamine 3000 (Thermo Fisher Scientific) (Figure 1A).

**Table 3.**
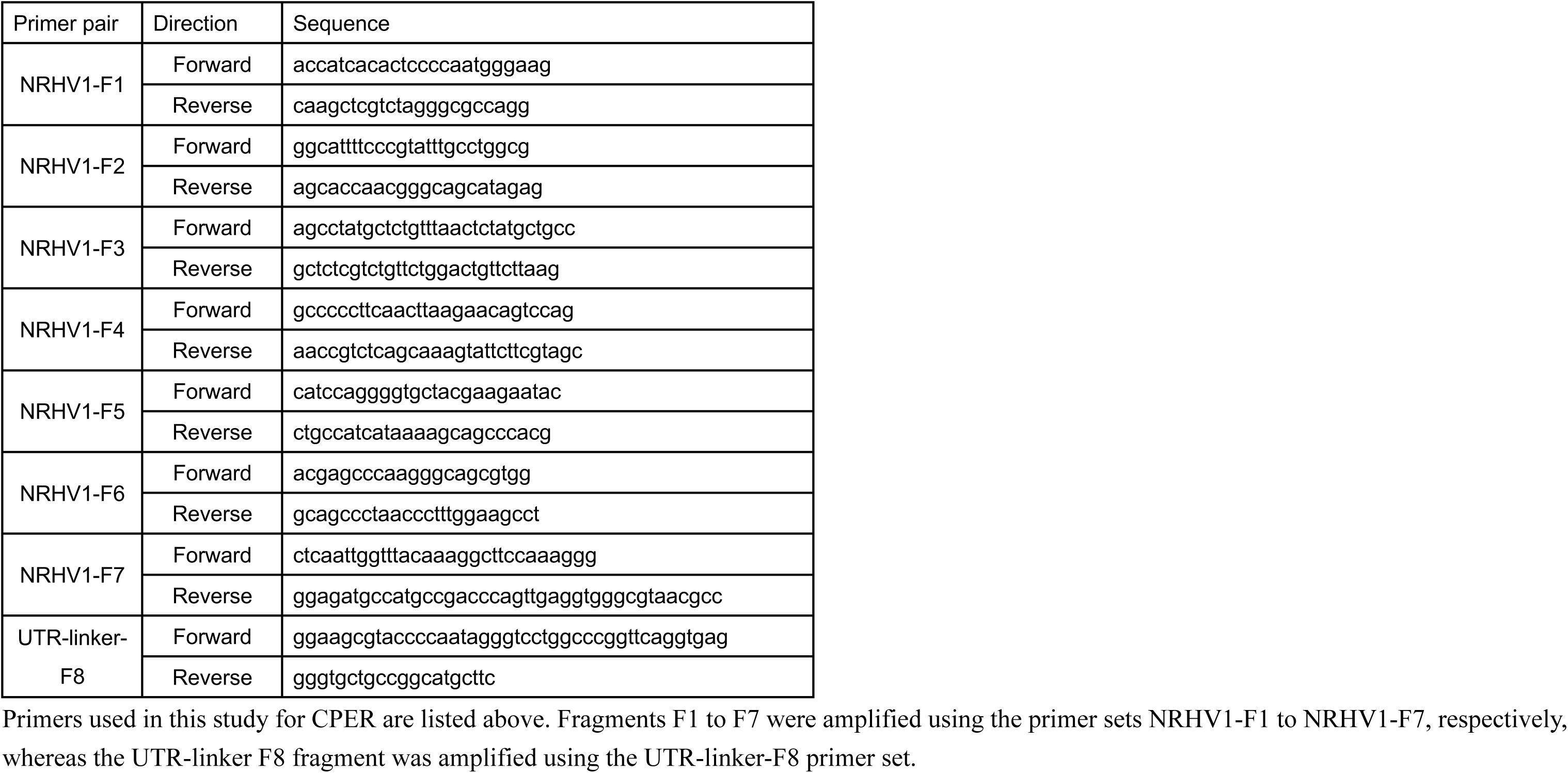
List of primers used for CPER.

### Antibodies and reagents

A rabbit polyclonal antibody against the NRHV1 core protein was commercially developed by immunization with the synthetic peptide SAAYPVRDPRRK (Scrum Inc., Japan). Rabbit polyclonal antibodies against NRHV1 NS5A were also commercially developed using synthetic peptides corresponding to two regions of the protein: clone #2 (KPLPKTKPTDPDTF) and clone #3 (YETSNDHVPKEDSW) (Scrum Inc., Japan). Clone #2 was used for Western blotting of NS5A derived from rNRHV1 HiBiT Δ378–391, as the epitope recognized by clone #3 is partially deleted in this construct. Anti-GAPDH (MBL, M171-3), anti-HiBiT (Promega, N7200) and anti-dsRNA clone K2 (Nordic MUbio, 10030005) antibodies were purchased. The compounds pibrentasvir and daclatasvir were purchased from Selleck.

### Extraction and quantitation of NRHV1 RNA from infected cells and culture supernatants

Total RNA was extracted from cells using an RNeasy Mini Kit (Qiagen, Hilden, Germany). Viral RNA in the supernatant was extracted using a QIAamp Viral RNA Mini Kit (Qiagen). The RNA samples were subsequently reverse transcribed to cDNA using ReverTra Ace qPCR RT Master Mix (TOYOBO). To detect viral RNA, qRT–PCR was performed using SYBR Green qPCR Master Mix (Applied Biosystems) and a QuantStudio 3 Real-Time PCR System (Applied Biosystems). The primer sets for the detection of viral RNA and the internal control were as follows: NRHV1 NS3: GGCAAGTTACCACCAGTGCT (sense) and ACTGGGATTAAGCACGAGGAC (antisense); GAPDH: ATGACTCTACCCACGGCAAG (sense) and CTGGAAGATGGTGATGGGTT (antisense). The level of viral RNA target relative to that of GAPDH mRNA was calculated as the delta Ct [ΔCt = Ct (NS3) − Ct (GAPDH)]. The amount of NRHV1 RNA in the supernatant was quantified by developing a standard curve using known copy numbers of in vitro-transcribed NRHV1 RNA.

### Immunofluorescence assay

Cells on glass slides were rinsed once with PBS, fixed with 4% paraformaldehyde for 10 min, and rinsed three times with PBS. The cells were then permeabilized in PBS supplemented with 0.1% saponin and 5% FBS at room temperature for 1 h. Immunofluorescence detection was performed using primary antibodies against NRHV1 NS5A (1:2000 dilution), HiBiT (1:1000 dilution), and dsRNA clone K2 (1:400 dilution) and appropriate secondary antibodies (IgG-Alexa488 or IgG- or IgM-Alexa594, Invitrogen), all of which were diluted in buffer containing PBS, 1% bovine serum albumin, and 0.1% saponin. Nuclei were stained with 4’,6-diamidino-2-phenylindole (DAPI; Abcam). Fluorescence images were obtained using a BIOREVO BZ-9000 (KEYENCE) or an A1R HD25 (Nikon).

### Western blotting

Cells were lysed on ice in lysis buffer (50 mM Tris-HCl (pH 7.5), 150 mM NaCl, 1% NP-40, and 1% glycerol) supplemented with a protease inhibitor cocktail (Nacalai Tesque). The lysates were boiled with loading buffer and subjected to 5 to 20% gradient SDS–PAGE. The proteins were transferred to polyvinylidene difluoride membranes (Millipore) using a semidry transfer blotter (ATTO). The membrane was incubated with Blocking One (Nacalai Tesque), treated with the appropriate antibodies and then washed with TBS containing 0.02% Tween-20. The immune complexes were visualized using SuperSignal West Femto maximum sensitivity substrate (Thermo Fisher Scientific) and detected by a FUSION chemiluminescence imaging system (M&S Instruments, Inc.).

### Phos-tag SDS–PAGE

Protein samples were separated on a 7.5% acrylamide gel supplemented with 50 µmol/L Phos-tag acrylamide (FUJIFILM Wako Chemicals) and 0.1 mmol/L MnCl₂. The gel composition was otherwise identical to that used for standard SDS–PAGE. After electrophoresis, the gels were incubated in transfer buffer containing 1 mmol/L EDTA for 20 min to chelate Mn²⁺, followed by incubation in transfer buffer without EDTA for more than 30 min before being transferred to PVDF membranes. Immunoblotting was then performed as described above.

### AlphaFold2 prediction

The amino acid sequences of E2, NS3, and NS5A of HCV and NRHV1 were input into ColabFold v1.5.5, which runs a simplified version of AlphaFold2 (https://colab.research.google.com/github/sokrypton/ColabFold/blob/main/AlphaFold2.ipynb) [35, 36]. The rank 1 models were visualized using PyMOL v2.5.8 (Schrodinger, L. & DeLano, W., 2020. PyMOL, available at http://www.pymol.org/pymol).

### HHpred analysis

The sequence and structural homology of NRHV1 NS5A domain I were assessed using HHpred (https://toolkit.tuebingen.mpg.de/tools/hhpred) [37, 38], a web-based tool that compares profile hidden Markov models (HMMs) to detect distant evolutionary relationships with higher sensitivity than pairwise sequence alignment methods. The full-length NRHV1 NS5A amino acid sequence was submitted as a query, and the resulting alignment indicated high similarity between residues 36–181 of NRHV1 NS5A and domain I of HCV NS5A (PDB ID: 3FQM).

### HiBiT and NanoLuc assays

HiBiT and NanoLuc activities were measured using Nano-Glo HiBiT lytic detection system and Nano-Glo luciferase assay system (Promega), respectively, in accordance with the protocol provided by the manufacturer.

### Neutralization assay

McA1.8 cells were seeded in 24-well plates and incubated overnight. rNRHV1 HiBiT was incubated for 1 h at 37°C with serum obtained from SD rats either 26 weeks post-infection (wpi) with NRHV1 or from naïve controls. The McA1.8 cells were then inoculated with the virus-serum suspension at 5 copies/cell. After a 4 h incubation, the cells were washed, and fresh medium was added. Luciferase activity was determined at 48 h post infection (hpi). Neutralizing activity was calculated relative to that of cells inoculated with the virus and incubated with naïve serum. To compare the timing of neutralizing antibody activity, serum samples were either preincubated with the virus before infection (preinfection) or added to the cells 4 h after inoculation.

### Animal experiments

Male, 5-week-old NOD.CB17-Prkdcscid/J (NOD–SCID) mice were purchased from Jackson Laboratory and allowed to acclimate for one week. For NRHV1 infection, mice were immobilized in a restrainer and then intravenously inoculated via the tail vein with culture supernatant containing recombinant NRHV1. Mock-infected mice received the same volume of culture supernatant from naïve cells. To monitor viremia, 10 µl of blood was collected from the tail vein at 2-week intervals post-infection and diluted to 140 µl with PBS. For the quantification of NRHV1 RNA, total RNA was extracted from mouse blood using a QIAamp Viral RNA Mini kit (QIAGEN), after which first-strand cDNA was synthesized using ReverTra Ace qPCR RT Master Mix (TOYOBO). To detect viral RNA, qRT–PCR targeting the NS3 region of NRHV1 was performed using a TaqMan probe and THUNDERBIRD qPCR Mix (TOYOBO). The TaqMan probe was labeled with 6-carboxyfluorescein (6-FAM) as the reporter dye and TAMRA as the quencher.

The NRHV1 NS3-specific primer and probe sequences used were as follows:

- Sense: TACATGGCTAAGCAATACGG;
- Anti-sense: AAGCGCAGCACCAATTCC; and
- Probe: 6FAM-CTCACGTACATGACGTACGGCATG-TAMRA.

The total RNA from tissue was purified with an RNeasy Protect Mini Kit (QIAGEN) and subjected to qRT–PCR as described for the infected cell samples. The HiBiT assay samples from organs were homogenized using a Handy Sonic in lysis buffer (100 mM Tris-HCl, 2 mM EDTA, 0.1% Triton X-100), and the supernatants obtained after centrifugation at 14000 ×g for 30 min at 4°C were measured using a BCA protein kit (Nacalai Tesque) to determine the protein concentration, after which 10 µg of protein from the supernatants was used for the HiBiT assay. For histological analysis, the mice were anesthetized and perfused with PBS, followed by 4% paraformaldehyde (PFA). The liver was then fixed in 10% neutralized buffered formalin solution.

All animal procedures were conducted in accordance with the Guidelines for the Care and Use of Laboratory Animals and were approved by the Institutional Committee of Laboratory Animal Experimentation of the University of Yamanashi.

### Immunohistochemistry and in situ hybridization of liver tissues

Paraffin-embedded liver sections (4 μm) were deparaffinized and processed for hematoxylin and eosin staining, immunohistochemistry, or in situ hybridization. For immunohistochemistry, the sections were washed with PBS, and antigens were retrieved by incubating the sections in pH 9 antigen retrieval buffer (Agilent DAKO) at 99°C for 15 minutes. After being washed with PBS, the sections were treated with Bloxall (Vector Laboratories) at room temperature for 15 min. Subsequently, the sections were blocked using a Histofine Mousestain Kit (Nichirei Biosciences, Tokyo, Japan), incubated with an anti-HiBiT antibody (1:250) at 4°C overnight, after which staining was visualized using a Histofine DAB substrate kit (Nichirei Biosciences). In situ hybridization was carried out using an RNAscope 2.5 HD Detection Reagent RED kit (ACD, Farmington, UT) and a probe targeting the helicase sequence (V-RHV-Rn1-NS3; cat. no. 504521) in accordance with the manufacturer’s protocol. Nuclei were counterstained with hematoxylin.

### Sequencing of viral RNA

Viral genome libraries were prepared using a NEBNext^®^ Ultra II RNA Library Prep Kit and XGen™ hybridization capture of Illumina^®^ Nextera™ DNA libraries. Sequencing was carried out on the Illumina iSeq 100 platform, and the resulting reads were analyzed using CLC Genomics Workbench software to identify viral mutations.

### Statistics

The data used for the statistical analyses are expressed as means ± standard deviations. The significance of differences between two groups was determined by Student’s t test. Significant differences among multiple groups were estimated by the Tukey–Kramer test. (*, p < 0.05; **, p < 0.01).

## Acknowledgments

We would like to thank Masako Mori for secretarial work and Midori Yoda, Akiko Yazaki and Keiko Ohmaki for technical assistance. This work was supported by the Japan Science and Technology Agency (JST) Moonshot R&D under grant number JPMJMS2025, by the Japan Agency for Medical Research and Development under Grant Number JP24fk0210109 and 25fk021016, by the Japan Society for the Promotion of Science KAKENHI Grant Number JP25K02498 and JP25K10374, and by a scholarship donation from Yakult Co., Ltd.

---

**Supplementary Figure 1.**
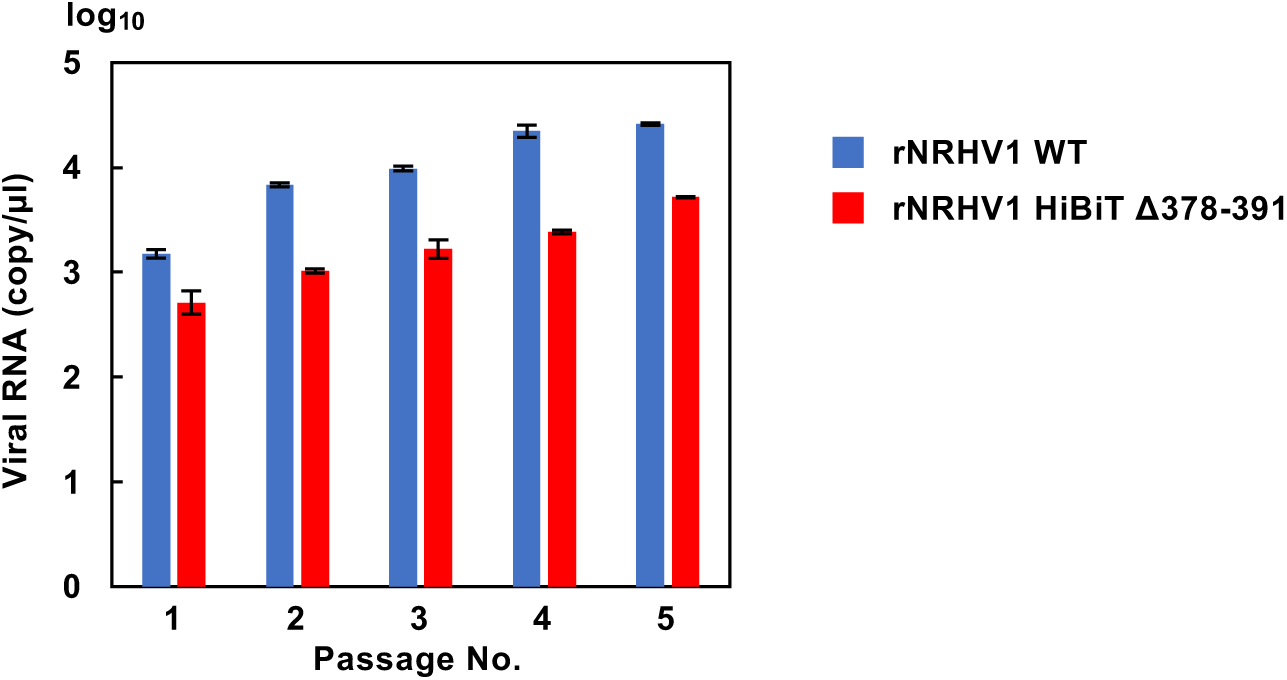
Generation of rNRHV1 virus stock. Culture supernatants collected from cells transfected with CPER products at 6 days post-transfection (dpt) were designated as passage 0 (P0). P0 supernatant was inoculated into naïve McA1.8 cells, and the resulting supernatant collected at 96 hours post-infection (hpi) was designated as P1. This passaging procedure was repeated sequentially to generate supernatants designated as P2 through P5. Viral RNA copy numbers in the supernatants were quantified by qRT-PCR.

**Supplementary Figure 2.**
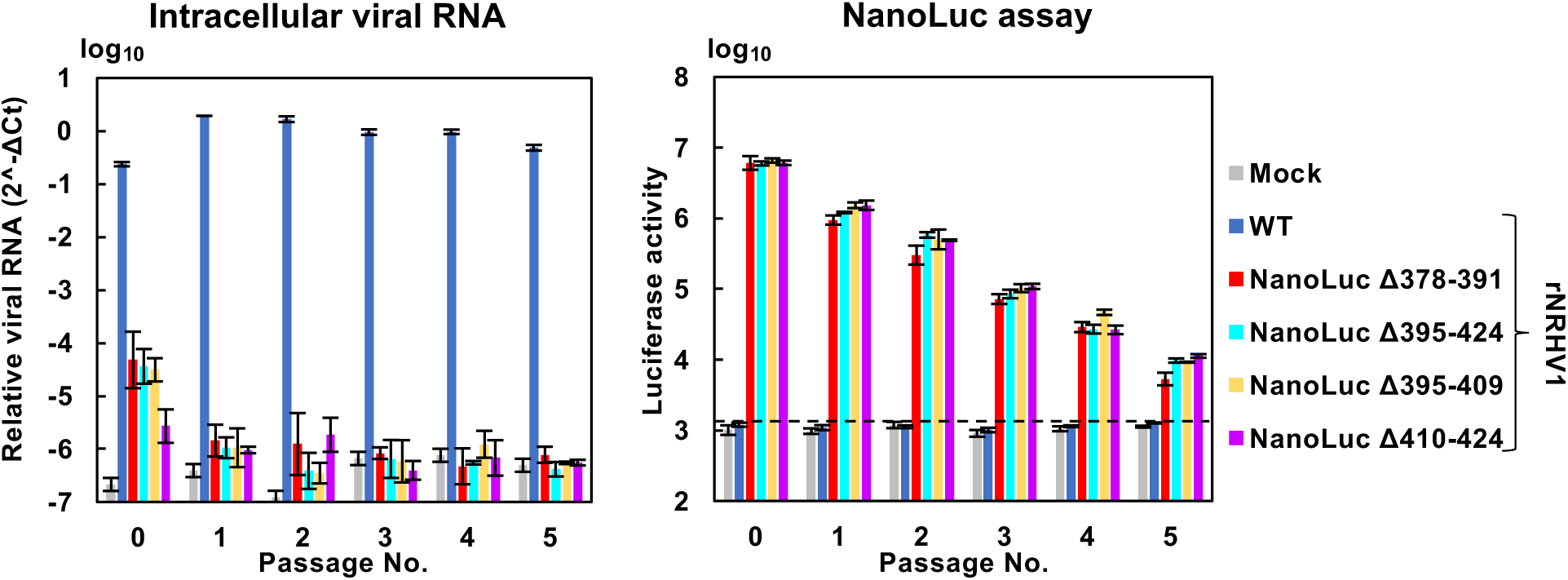
Stability of rNRHV1 expressing NanoLuc *in vitro*. Supernatants were collected from cells transfected with CPER-derived viral preparations at 6 dpt and designated as passage 0 (P0). P0 supernatants were then inoculated into naïve McA1.8 cells, and the resulting supernatants collected at 96 hpi were designated as P1. This passaging procedure was repeated sequentially to generate P2 through P5. Subsequently, 100 µl of each passage supernatant (P0–P5) was inoculated into naïve McA1.8 cells. Intracellular NRHV1 RNA levels and luciferase activity were assessed at 96 hpi.

**Supplementary Figure 3.**
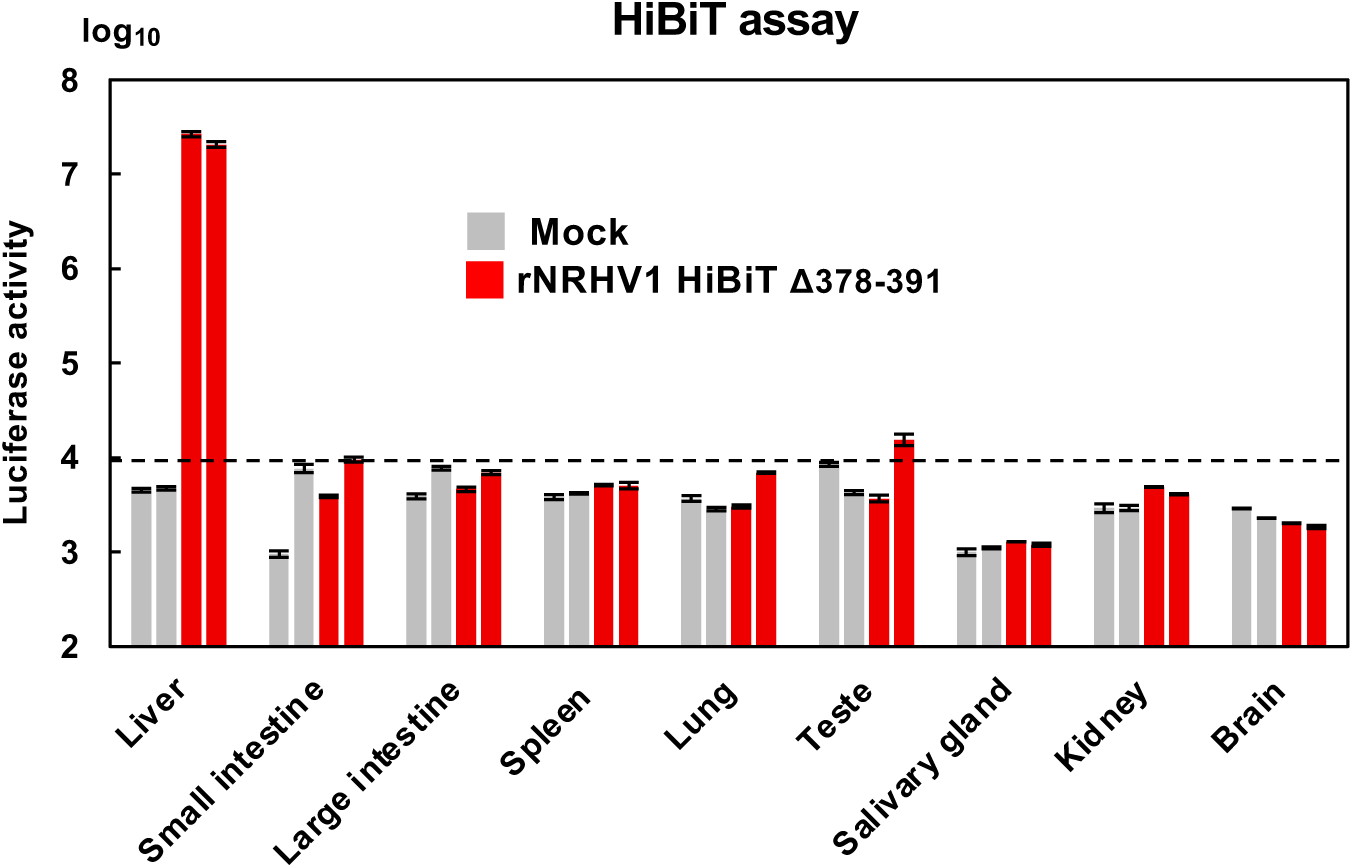
HiBiT activity in various organs of mice infected with rNRHV1 HiBiT Δ378–391. HiBiT activity in 10 µg of tissue lysate was measured. Individual organs were collected from rNRHV1 HiBiT Δ378–391–infected mice (mouse No. 1 and No. 3) and from a mock-infected control mouse.

**Supplementary Figure 4.**
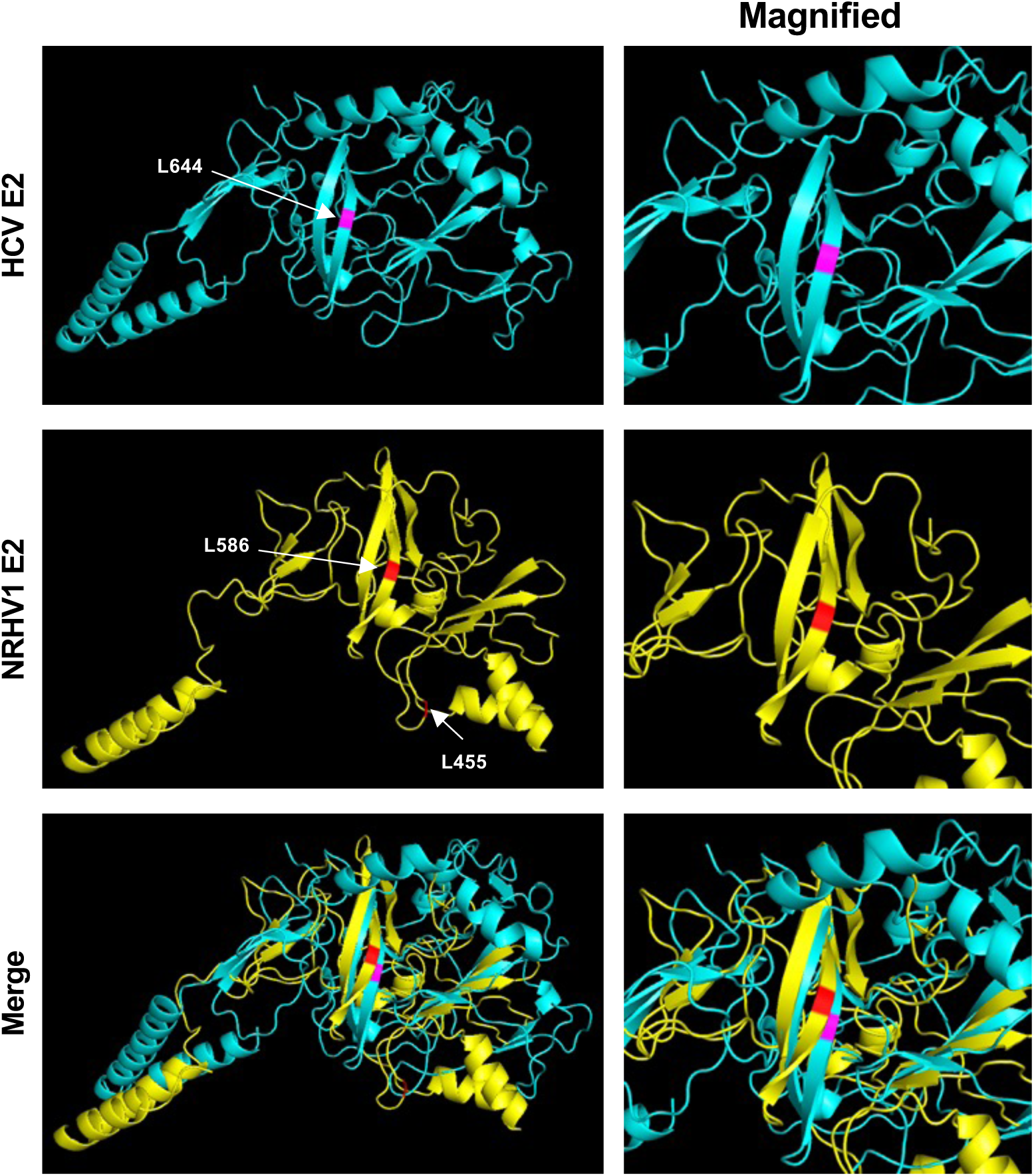
Structural comparison between HCV E2 and NRHV1 E2. The 3D structures of HCV E2 (blue) and NRHV1 E2 (yellow) were predicted using AlphaFold2.

**Supplementary Figure 5.**
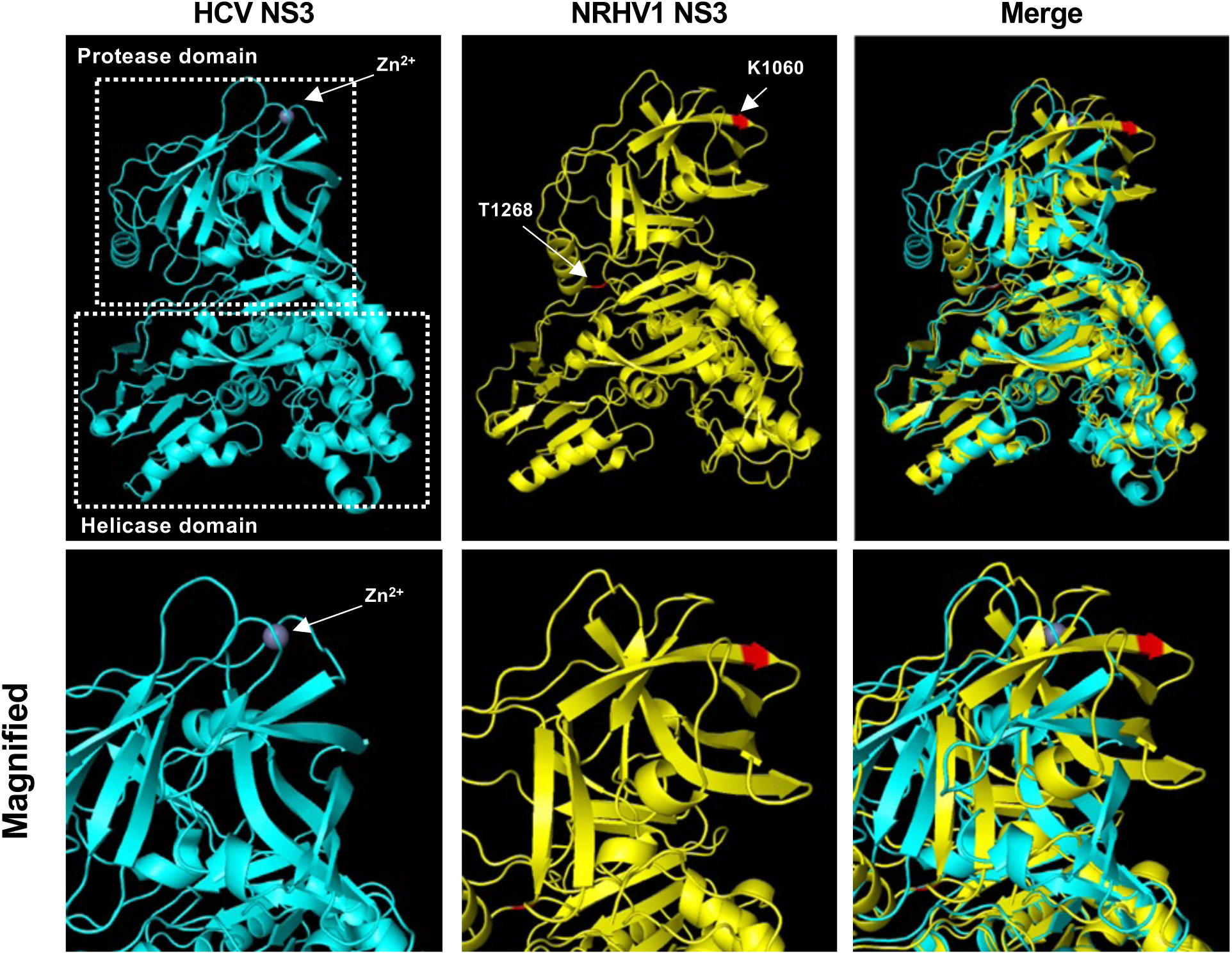
Structural alignment of HCV and NRHV1 NS3. Ribbon diagrams show the crystal structure of HCV NS3 (PDB ID: 3O8B, blue) and the predicted structure of NRHV1 NS3 (yellow).

